# Impaired active myocardial relaxation is the earliest sign of anthracycline cardiotoxicity and reflects selective injury of the cardiomyocyte surface

**DOI:** 10.64898/2025.12.05.692508

**Authors:** Céline Guilbeau-Frugier, Clément Karsenty, Ramona Ghenghea, Marie Cauquil, Véronique Lachaize, Nicolas Pataluch, Olivier Lairez, Etienne Dague, Childerick Severac, Jean-Michel Sénard, Céline Galés

## Abstract

**Aims:** Anthracycline-induced cardiotoxicity is a leading cause of heart failure (HF) in cancer survivors and is usually detected only once systolic dysfunction is established, when myocardial injury may already be advanced. Diastolic dysfunction may occur earlier, but its onset and cellular basis are unknown. We investigated the temporal emergence of diastolic dysfunction during doxorubicin (DOX) exposure and its link to the selective vulnerability of crest/subsarcolemmal mitochondria (SSM) at the cardiomyocyte (CM) surface.

**Methods and results:** Using a clinically relevant chronic DOX protocol in adult mice, we performed longitudinal cardiac assessment from the first exposure, combining conventional echocardiography, tissue Doppler, and global longitudinal strain (GLS). Structural and molecular correlates were characterized by atomic force microscopy, transmission electron microscopy, and 3D-quantitative imaging of native cardiac tissue, spatially resolving the CM surface (SSM) from the interior (interfibrillar mitochondria, IFM) compartment.

DOX impaired active myocardial relaxation from the first dose with prolonged isovolumic relaxation time and reduced mitral annular e′, whereas passive filling (transmitral filling, left atrial size) was preserved. Ejection fraction remained normal, with an early reduction in GLS. This early relaxation defect coincided with selective remodeling of the surface crest/SSM architecture, evident as crest flattening and progressive SSM loss and with depletion of the crest/SSM determinants Ephrin-B1 and Claudin-5, whereas IFM were affected only later, upon cumulative exposure. Surface SSM injury was accompanied by early, spatially ordered PINK1/Parkin activation, engaging SSM before IFM, and by loss of the sarcolemmal Na⁺/Ca²⁺ exchanger NCX1, likely impairing diastolic Ca^2+^ removal. In CM-specific Ephrin-B1-deficient mice, which lack mature crest/SSM, DOX precipitated an accelerated transition to HF with reduced ejection fraction.

**Conclusions:** Impaired active relaxation is the earliest functional hallmark of anthracycline cardiotoxicity, underpinned by selective injury of the surface crest/SSM compartment and its Na⁺/Ca²⁺ exchanger NCX1, while IFM and passive filling are initially spared. Diastolic assessment from the first anthracycline cycles may improve early detection and prevention of HF progression.

## METHODS

### Ethical approval

All animal experiments were performed in accordance with the European Directive 2010/63/EU for the protection of animals used for scientific purposes, and approved by the French Ministry of Research and the local institutional ethics committee (CEEA-122, Protocol No. CEEA 122-2014-33). All procedures complied with institutional and national guidelines for the care and use of laboratory animals.

### Animal models

All experiments were conducted on adult, 2-month-old male C57BL/6J OlaHsd mice (purchased from Envigo, Huntingdon, United-Kingdom). Tamoxifen-inducible cardiomyocyte-specific *Efnb1* conditional knock-out mice (cKO) and their WT littermate controls have been previously described^1^. All animals were housed in the animal facility of the “UMS06-Centre régional d’exploration fonctionnelle et de resources expérimentales-CREFRE (Toulouse, France)” under specific-pathogen free (SPF) conditions and following institutional animal use and care guidelines. Mice were maintained in conventional cages under controlled temperature and humidity, with a 12-hour light/dark cycle and free access to food and water. Sample size estimation was based on previous studies from our group. For excision of the beating heart, mice were deeply anesthetized with an intraperitoneal injection of 50 mg/kg Dolethal (Vetoquinol, Lure, France), and 0.1 mg/kg buprenorphine was topically applied to the subcostal incision site to minimize local nociceptive responses. The diaphragm was then rapidly incised, the rib cage lifted and the beating-heart was promptly excised using curved forceps and small scissors at the base. Euthanasia was subsequently completed by immediate cervical dislocation.

### Purification of adult cardiomyocytes from mouse hearts

Adult CMs from the left ventricle were freshly isolated from beating mouse hearts using a retrograde Langendorff perfusion-based enzymatic digestion protocol, as previously described^2,3^.

### Doxorobucin treatments

For *in vivo* DOX-treatments, two-month-old mice received intraperitoneal injections of doxorubicin (DOX, 5 mg/kg body weight; Sigma-Aldrich, D1515) once weekly for up to 5 weeks. The impact of DOX on CM architecture was evaluated after cumulative doses of 5, 20, and 25 mg/kg. Hearts were harvested 4 days after DOX injection. Long-term DOX-induced cardiotoxicity was assessed in a separate cohort, in which hearts were collected 15 weeks after the final injection (cumulative dose: 25 mg/kg + 15 weeks).

For *in vitro* DOX treatments, immediately after adhesion, freshly isolated adult CMs were exposed or not (control) to increasing concentrations of DOX (1, 1.5, 2, 2.5 or 5 µM) in CM culture medium (Minimum Essential Medium-MEM, 5% Fetal Calf Serum, 100 U.mL^-1^ Penicilin, 2 mM.L-1 Glutamin) supplemented with 10 mM 2,3-butanedione monoxime (BDM) to prevent contraction. Cells were incubated with DOX for 5 minutes at 37°C in a humidified 5% CO₂ atmosphere. Following treatment, cells were washed and transferred to CM culture medium supplemented with BDM prior to AFM measurements.

### Echocardiography, speckle tracking strain analysis, and Doppler imaging

Transthoracic echocardiography was performed under light anesthesia with 1% isoflurane in 0.5% oxygen, using a Vevo® 2100 system (VisualSonics) equipped with a 40 MHz MS550D transducer. All measurements were obtained by an examiner blinded to the animal groups, and data were averaged over three consecutive cardiac cycles.

Left ventricular ejection fraction (LVEF) was assessed in parasternal long-axis B-mode, measured at least three times per animal. LV wall thickness and internal diameters were quantified from M-mode images at the level of the papillary muscles in both long- and short-axis views. LVEF (%) was calculated from B-mode images by tracing end-diastolic and end-systolic endocardial borders to estimate LV volumes using the formula:EF (%) = [(EDV − ESV) / EDV] × 100.

Global longitudinal strain was derived from parasternal long-axis views using speckle-tracking-based analysis to assess global systolic performance.

Diastolic function was evaluated by pulsed-wave Doppler of mitral inflow from the apical four-chamber view, measuring early (E) and late (A) transmitral flow velocities, isovolumic relaxation time (IVRT), and calculating the E/A ratio. Tissue Doppler imaging (TDI) was used to assess early (e′) and late (a′) diastolic mitral annular velocities, and the E/e′ ratio was derived.

### Atomic Force Microscopy (AFM) Force-Volume mapping on living cardiomyocytes

AFM measurements were performed on freshly isolated adult CMs from control or DOXO-treated mice. All measurements were obtained by an examiner blinded to the animal groups. Following isolation, dissociated adult CMs were plated onto culture dishes pre-coated with laminin-2 (10 µg/mL) at 37°C under 5 % CO_2_. After 20 minutes to allow cell attachment, the plating medium was replaced with a CM culture medium (Minimum Essential Medium-MEM, 5% Fetal Calf Serum, 100 U.mL^-1^ Penicilin, 2 mM.L^-1^ Glutamin) supplemented with 10 mM 2,3-butanedione monoxime (BDM) to prevent spontaneous contraction during AFM acquisition and thereby better preserve crest/SSM at the CM surface, which are highly sensitive to contraction in isolated CMs. This contraction-free acquisition is required for reliable AFM mapping.

AFM measurements were conducted using a NanoWizard II atomic force microscope (JPK Instruments, Berlin, Germany) equipped with MLCT cantilevers (Bruker, Santa Barbara, CA, USA), composed of silicon nitride (Si₃N₄) and characterized by a pyramidal tip geometry with an average tip radius of curvature of 35 nm. All recordings were performed in the CM culture medium at 37°C under continuous perfusion and 5% CO_2_, using the PetriDishHeater™ perfusion system and the integrated temperature control unit of the NanoWizard II setup.

Height images and elasticity maps were acquired in Force-Volume mode, as previously described^2^, at a resolution of 64 × 64 force curves per map over a scan area of 100 µm². For each cantilever, both deflection sensitivity and spring constant were calibrated prior to use. The spring constant was determined using the thermal noise method, and ranged between 0.101 and 0.120 N/m. To preserve CM plasma membrane integrity, the maximum applied force was limited to 3 nN. Force-distance curves were converted to indentation curves after calibration, and the Hertz model was applied to calculate the apparent Young’s modulus as a measure of cell surface stiffness. Force curves were extracted from the elasticity maps for further quantitative analysis.

### Transmission electron microscopy (TEM) and quantitative analysis

Heart samples were specifically processed to preserve CM surface crests as previously described^1,3^. Briefly, immediately after excision, hearts were rinsed in cold PBS and carefully sectioned into ∼1 mm³ biopsies on a glass slide placed over an ice bed. Samples were then fixed in 2% glutaraldehyde in Sorensen’s buffer at 4 °C and processed for TEM. Each biopsy was embedded in a separate resin block. The methodology for quantitative analysis of crest/SSM on TEM images has been extensively described previously^1,3^.

### Formalin-fixed paraffin-embedded (FFPE) heart sections

Immediately after excision, beating hearts were rinsed three times in cold PBS, fixed in 4% formaldehyde for 48h, embedded in paraffin, and sectioned in 5 μm transverse sections for subsequent analysis.

### 2D quantification of CM area

For *in situ* quantification of CM surface area, FFPE heart sections were deparaffinized and stained for 1 hour at room temperature with OG488-conjugated-Wheat Germ Agglutinin (WGA) (Invitrogen, W6748; 10 µg/mL in HBSS) to enable precise delineation of the CM surface, in the presence of the nuclear probe 4,6-diamidino-2-phenylindole (DAPI) (Sigma-Aldrich, 32670, 1:1000°). CM area was measured on transverse heart cross-sections at the endocardial level (left ventricle) by manually tracing cell contours on whole-heart images acquired with a digital *NanoZoomer* slide scanner (Hamamatsu) and analyzed using Zen 2011 software (Carl Zeiss). The experimenter was blinded to group allocation.

### Quantification of cardiac fibrosis

FFPE heart sections were stained using a Masson’s Trichrome Stain Kit (Abcam, ab150686). Fibrosis was quantified on trichrome-stained heart sections using a custom ImageJ macro applied to whole-heart images acquired with a digital *NanoZoomer* slide scanner (Hamamatsu). The experimenter was blinded to the mouse genotype.

### Fluorescent immunolabeling and confocal imaging of fresh cardiac biopsies

Fresh cardiac biopsies were immunolabeled and imaged using the 3D-NaissI (3D-Native Tissue Imaging) method, as previously described^4^. Briefly, fresh biopsies were incubated in HBSS with the primary antibody for 2 days at 4 °C under constant rotary agitation, then washed and incubated with the fluorescent secondary antibody, protected from light, for a further 2 days under the same conditions. Biopsies were then imaged on an inverted ZEISS LSM 900 Axio Observer.Z1/7 confocal microscope using a ×63 oil-immersion objective (NA 1.4) at a maximum resolution of 1437 × 1437 pixels (pixel scaling 0.071 × 0.071 µm). Three-dimensional stacks (10 µm thickness) were acquired with a z-step of 1 µm, starting below the tissue surface to exclude the most superficial sections (first ∼4 µm) and avoid surface-related artefacts, as previously described⁴. Emission filter ranges, detector gains, and pixel dwell time (0.54 µs) were set as detailed in Pataluch et al.^4^. Images were visualized with Zen Blue software (Carl Zeiss, Germany). Acquisition parameters were kept identical across conditions for a given primary antibody, but were adjusted between different antibodies. The images shown are representative of several independent acquisitions.

### Spatially resolved quantification of surface (SSM) and interior (IFM) protein expression in cardiomyocytes within fresh cardiac biopsies

To separately quantify protein expression at the CM surface (reflecting SSM), and within the CM interior (reflecting IFM), 2D fluorescence images acquired by 3D-NAissI (transverse sections of left ventricular myocardium) were analyzed with a custom Fiji/ImageJ macro enabling automated, cell-by-cell quantification.

For each biopsy, three 2D image stacks were analyzed (top, middle and bottom of the sample), each capturing the same CMs (three stacks per CM); the most superficial stacks (first 4 µm) were excluded as previously described^4^. After import via Bio-Formats, CMs were segmented with Cellpose, custom-trained to delineate individual CMs from the inner edge of the wheat-germ agglutinin (WGA) surface staining, generating a 16-bit CM label image. Within each segmented CM, a surface mask was defined as a 10-pixel-thick band beneath the WGA signal (erosion 1, dilation 10), reflecting the surface SSM compartment; the remaining internal area, after exclusion of nuclei (DAPI staining), defined the intracellular mask, reflecting the IFM compartment.

Protein expression was quantified from the number of fluorescent pixels above threshold rather than from mean fluorescence intensity, because the tissue and inter-experiment variability inherent to the 3D-NaissI method precludes reliable comparison of absolute fluorescence intensities^4^. For each protein, the binary threshold was set on the control condition of each individual experiment and applied identically to all conditions. Within each CM, the number of threshold fluorescent pixels was measured separately in the surface and intracellular masks (MorphoLibJ 2D region analysis) and normalized to the area of the corresponding mask, yielding a surface (SSM) and an intracellular (IFM) protein expression value per CM. Values were computed across the three sections per biopsy (3 mice; 5 biopsies/mouse; 3 sections/biopsy; ∼20–70 CMs/mouse), exported as CSV, and expressed as a percentage of the control mean.

### Data analysis-statistics

The n number for each experiment and analysis is stated in each figure legend. The normality was tested using the Shapiro-Wilk normality test. Results are reported as mean ± SD. An unpaired Student t-test for parametric variables was used to compare two groups. One-way ANOVA with Tukey post-hoc test was used for multiple group comparisons (> 2 groups). The level of significance was assigned to statistics in accordance with their p values: p ≤ 0.05 *; p ≤ 0.01 **; p ≤ 0.001 ***; p ≤ 0.0001 ****). All graphs were generated using GraphPad Prism v10.1.2 (GraphPad Inc, San Diego, California).

## INTRODUCTION

Doxorubicin (DOX), a widely used anthracycline, remains among the most effective chemotherapeutic agents for adult and pediatric cancers including breast cancer, leukemia and lymphoma^5^. Its introduction has substantially improved cancer survival. However, its clinical use is limited, by cumulative, dose-dependent cardiotoxicity which may emerge months to years after treatment and markedly raises the risk of heart failure (HF)^6,7^. This delayed toxicity represents a major long-term burden for cancer survivors and has contributed to the development of dedicated cardio-oncology programs^8^.

Anthracycline cardiotoxicity is nevertheless still frequently identified only after systolic dysfunction has developed, when myocardial injury may already be advanced and only partly reversible. Several strategies have been explored to reduce DOX-induced cardiotoxicity (DIC), including dose limitation, co-administration of cardioprotective agents, and the use of less toxic anthracycline formulations or analogues^9–11^. However, these approaches may provide incomplete protection and, in some settings, interfere with anticancer treatment. Notably, emerging evidence suggests that DOX-induced cardiomyopathy may be at least partly reversible when detected at a subclinical stage and treated promptly^7,12^. Current surveillance of patients receiving DOX relies primarily on left ventricular ejection fraction (LVEF), global longitudinal strain (GLS), and circulating biomarkers, usually performed at baseline, selected time-points during treatment and across the first year^7^. Early increases in cardiac troponin predict subsequent ventricular dysfunction^13^, but do not, alone, justify altering therapy. Several prospective studies, including the largest to date, have reported early abnormalities in diastolic function during anthracycline therapy that may precede systolic dysfunction^14–17^. These observations support a model in which impaired relaxation represents an early stage of anthracycline cardiotoxicity, although the value of diastolic indices in this setting remains uncertain, in part because of their load dependence^18^. Although diastolic parameters are included in the comprehensive echocardiographic assessment recommended by current European Society of Cardiology (ESC) cardio-oncology guidelines^5^, they are not incorporated into the definitions or therapeutic thresholds of cancer therapy-related cardiac dysfunction and are not specifically monitored from the first treatment cycles. Moreover, the cellular and molecular basis of this early diastolic impairment remains currently unknown. There is therefore a need to identify early functional abnormalities and their underlying mechanisms from the initial cycles of anthracycline exposure.

Preclinical studies have established that DIC is a multifactorial process involving all major cardiac cell types^19^. In cardiomyocytes (CMs), DNA damage, topoisomerase IIβ inhibition, iron dysregulation, and excessive reactive oxygen species (ROS) generation have been implicated^11,20^. Among these, mitochondrial dysfunction is considered a central contributor, characterized by impaired bioenergetics, oxidative stress, and ultrastructural abnormalities^21^. However, the translational relevance of these findings remains limited. Most preclinical models rely on acute or high-dose DOX administration and focus on advanced HF^22,23^, making it difficult to distinguish initiating mechanisms from secondary consequences and poorly reproducing the progressive cumulative exposure in patients. Up to now, whether myocardial relaxation is impaired after the first DOX exposure, and which subcellular mechanisms underlie this very early response, remain unresolved.

We previously identified an architectural specialization of the lateral membrane in adult CMs from male mice and Human, consisting of arrays of regularly spaced bulges (referred to as “crests”), that contain a distinct population of subsarcolemmal mitochondria (SSM)^1–3^. *In situ*, crests form intermittent physical contacts with corresponding structures on adjacent CMs within the left ventricle myocardium. These crest-crest contacts involve the junctional protein Claudin-5 and may reinforce lateral tissue cohesion^3^. Recently, we have shown that crests/SSM mature lately after postnatal day 20, characterized by SSM swelling and associated with an atypical form of cardiac hypertrophy without myofibrillar addition^1^. This maturation is under the control of the lateral membrane protein Ephrin-B1^24^, a Claudin-5 partner^24^, and specifically drives the developmental gain in diastolic performance in males^1^, although the mechanisms linking crest/SSM organization to myocardial relaxation remain incompletely understood. Accordingly, CM-specific deletion of *Efnb1* disrupts crest/SSM maturation and induces selective diastolic dysfunction, an HFpEF phenotype that evolves with age toward HFrEF and 100% mortality by 13 months due to terminal HF¹. Crests/SSM are also selectively vulnerable to cardiac stress. They are among the earliest mitochondrial populations lost after myocardial ischemia^2^ and during pressure-overload-induced remodeling^3^, preceding interfibrillar mitochondrial (IFM) disorganization, T-tubule disruption, and CM death. These findings suggest that loss of crest/SSM at the CM surface may represent an initiating event in the progression of myocardial injury. Whether this spatially restricted mitochondrial vulnerability contributes to the earliest stages of anthracycline cardiotoxicity, and whether it provides a mechanistic substrate for early diastolic dysfunction, remain unknown.

In the present study, we used a preclinical model of chronic DOX protocol with longitudinal cardiac phenotyping beginning after the first drug exposure, combined with high-resolution structural imaging and spatially resolved molecular analyses. We investigated whether impaired active relaxation is detectable after a single DOX dose, defined the associated changes in SSM and IFM compartments, and examined whether disruption of the crest/SSM microdomain contributes to subsequent progression toward systolic dysfunction.

## RESULTS

### A chronic doxorubicin model recapitulates the clinical features of anthracycline cardiomyopathy and reveals very early functional and structural alterations

To recapitulate the clinical setting of cumulative anthracycline exposure, adult male C57BL/6J mice were administered DOX or saline (control) at 5 mg/kg once weekly for five consecutive weeks, reaching a total cumulative dose of 25 mg/kg (**Figure 1A**). Systolic function and myocardial/CM structure were assessed longitudinally at baseline (prior to the first DOX injection), 3 days after the first dose (5 mg/kg) to capture acute DOX effects, after 4 weeks (cumulative dose: 20 mg/kg), after 5 weeks (cumulative dose: 25 mg/kg), and 15 weeks after the final injection (25 mg/kg + 15 weeks) to assess long-term consequences.

**Figure 1.**
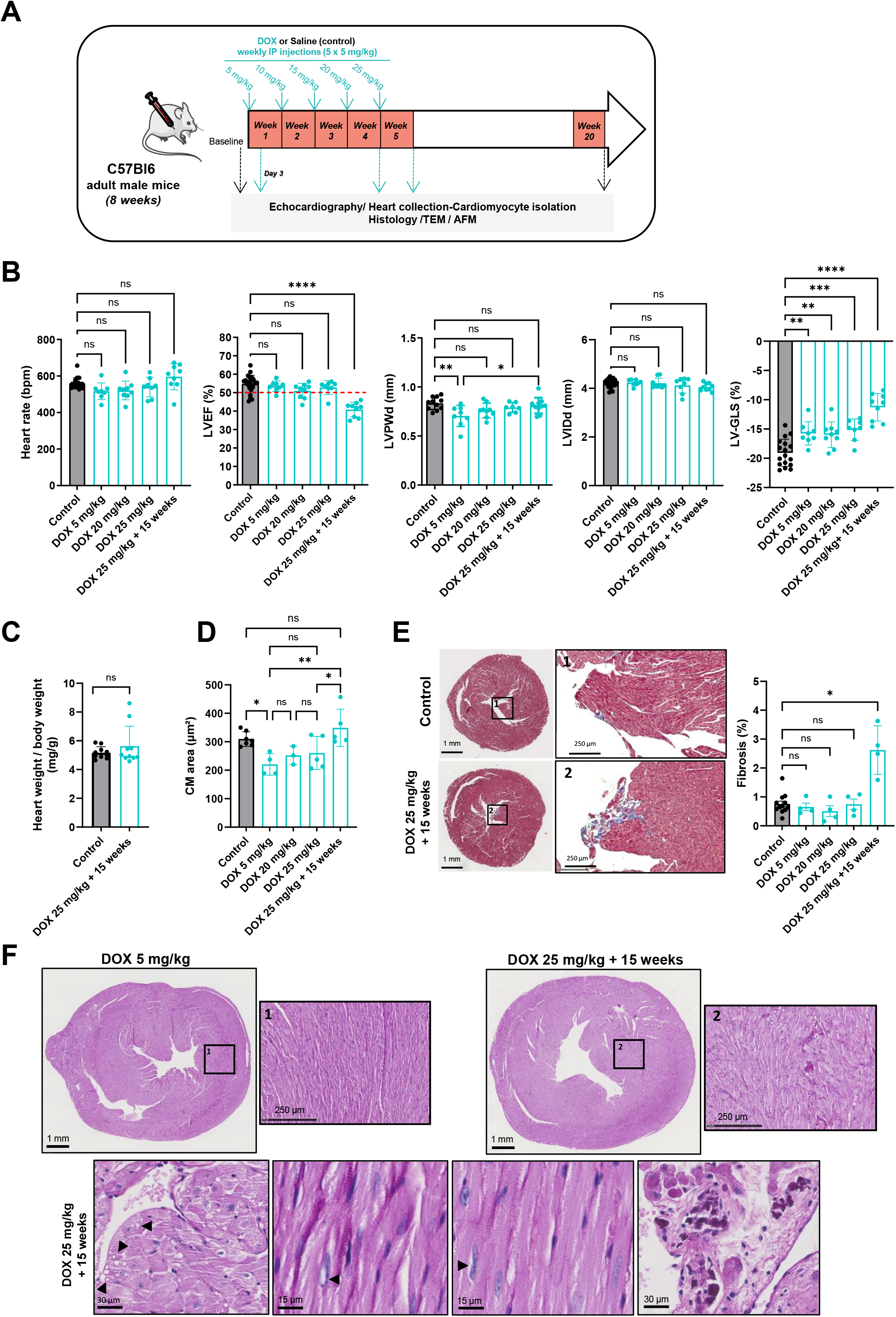
Early and progressive cardiomyocyte remodeling and functional decline in a murine model of chronic doxorubicin-induced cardiotoxicity. **(A)** Schematic timeline of the experimental protocol: adult male C57Bl/6J mice (8 weeks old) received weekly intraperitoneal doxorubicin (DOX; 5 mg/kg/week) or saline (control) for 5 weeks, (cumulative 25 mg/kg). Echocardiography was performed longitudinally at baseline, 3 days after the first dose (5 mg/kg), and after 4 (20 mg/kg), 5 (25 mg/kg) and 20 weeks (25 mg/kg + 15 weeks). Hearts were collected 3-4 days after the final injection (acute groups: 5, 20, 25 mg/kg) or 15 weeks later (chronic group) for structural and molecular analyses (histology, TEM, AFM). **(B)** Longitudinal echocardiography (control: n=20; DOX groups: n=8-10 mice/group; controls pooled from all saline-injected mice). Heart rate and left ventricular internal diameter in diastole (LVIDd) were unchanged. LVEF was preserved throughout the injection period and declined only at 15 weeks. LV posterior wall thickness in diastole (LVPWd) was transiently reduced after the first dose. LV global longitudinal strain (LV-GLS) was progressively impaired from the first dose (5 mg/kg). **(C)** Heart weight–to–body weight ratio at 15 (control: n=10; DOX 25 mg/kg: n=10), unchanged between groups. (**D**) Cardiomyocyte (CM) cross-sectional area from WGA-stained transverse heart sections (∼200 CMs/mouse; control: n=6 mice, DOX: n=3-5 mice/group), showing early reduction and later normalization. **(E)** Interstitial fibrosis from Masson’s trichrome-stained heart sections (control: n=4; DOX: n=4/group; controls pooled). *Right*: quantification. *Left*: representative images showing mild endocardial fibrosis in a late DOX-treated mouse. **(F)** Representative Periodic Acid-Schiff (PAS)-stained images of DOX-induced CM damage (DOX 25 mg/kg+15 weeks). From left to right (black arrows): cytoplasmic vacuolization, intranuclear light inclusion, perinuclear light halo, and cytoplasmic glycogen accumulation within areas of CM necrosis. Data are mean ± S.D. Statistics: one-way ANOVA with Tukey’s multiple comparisons test for all group-wise comparisons (B, D), Student’s *t* test for two-group comparisons (C), Kruskal-Walli’s test followed by Dunn’s multiple comparisons test to compare each DOX-treated groups to the control (E). * *P* <0.05, ** *P* <0.01, *** *P* <0.001, **** *P* <0.0001. ns: not significant.

Echocardiography showed preserved left ventricular ejection fraction (LVEF) throughout the DOX injection period, with a marked decline appearing only at the 15-week follow-up compared to controls (**Figure 1B**), mirroring the delayed clinical progression of DOX cardiomyopathy. No compensatory cardiac hypertrophy was detected at 15 weeks, as indicated by stable left ventricular posterior wall thickness in diastole (LVPWd) (**Figure 1B**), unchanged heart-to-body-weight ratio (**Figure 1C**) and CM cross-sectional area (**Figure 1D**). Only a mild but significant increase in endocardial fibrosis was observed (**Figure 1E**), indicating minimal structural remodeling at this stage. Histological analysis of 15-week cardiac tissue revealed typical features of DOX-induced injury, including CM intracellular vacuolization, intranuclear light inclusions, perinuclear clearing (light halo), and focal glycogen accumulation in limited areas of CM necrosis (**Figure 1F**).

In contrast, global longitudinal strain (GLS), a sensitive marker of subclinical cardiotoxicity, with a >15% relative reduction defining early injury per the 2022 ESC guidelines^5^, was impaired early. Compared with controls, in DOX-treated mice, LV-GLS became less negative from −18% at baseline to ∼−15% after a single DOX dose (5 mg/kg) and −10% at 15 weeks (**Figure 1B**), corresponding to relative reductions of ∼21% and ∼47%, respectively, both above the ESC threshold. This early GLS impairment was accompanied by a transient decrease in LVPWd immediately after the first DOX dose (5 mg/kg), which progressively normalized to control by 15 weeks (**Figure 1B**), paralleled by an early reduction in CM cross-sectional area that similarly normalized by 15 weeks (**Figure 1D**), consistent with early CM atrophy followed by late compensatory hypertrophic adaptation.

Thus, these findings validate this preclinical model of chronic anthracycline cardiotoxicity and demonstrate that DOX induces early functional and structural alterations from the first exposure, preceding any LVEF decline and the later development of HF.

### A single doxorubicin exposure is sufficient to impair active myocardial relaxation, while filling and atrial size are preserved

Because myocardial function was already perturbed after a single DOX dose, we next examined diastolic function in a separate cohort subjected to the same protocol (DOX 5 mg/kg once weekly; **Figure 2A**), and focused the diastolic analysis on two time points, 3 days after the first injection (5 mg/kg; acute effect), and after four weeks (cumulative 20 mg/kg). LV diastolic function was assessed by pulsed-wave and tissue Doppler, comparing DOX with time-matched saline controls at each dose.

**Figure 2.**
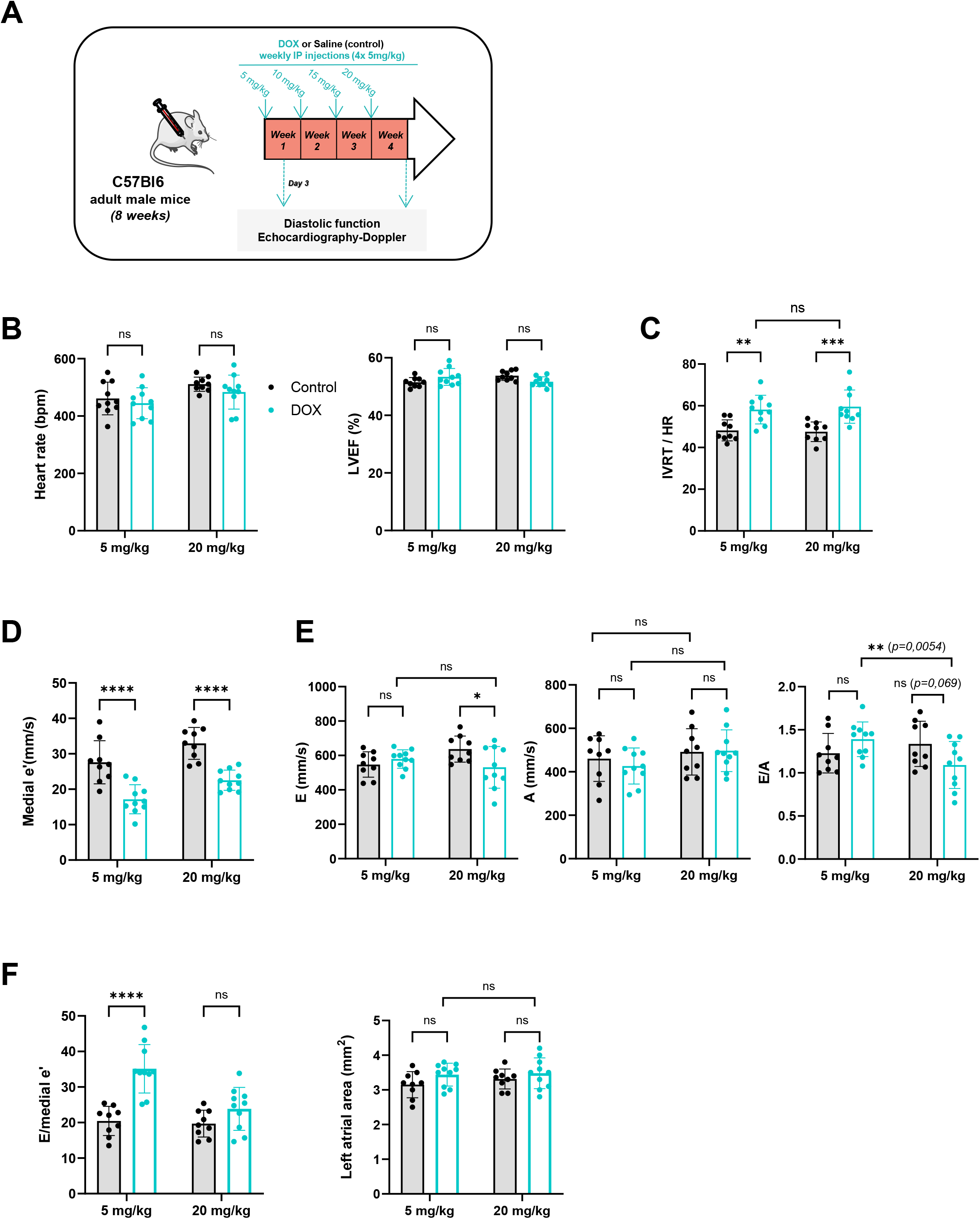
Early onset of diastolic dysfunction with preserved systolic function in adult mice following doxorubicin exposure. (A) Schematic representation of the *in vivo* protocol. Adult male C57BL/6J mice (8 weeks old) received weekly intraperitoneal injections of DOX (5 mg/kg) or saline (control) for up to 4 weeks (cumulative dose: 20 mg/kg). Longitudinal transthoracic echocardiography with pulsed-wave and tissue Doppler was performed 3 days after the first DOX injection (5 mg/kg) and one week after the fourth injection (20 mg/kg). (**B**) Heart rate and left ventricular ejection fraction (LVEF) were unchanged in DOX-treated mice at both doses, confirming preserved systolic function and excluding heart rate as a confounder. (**C**) Heart rate-corrected isovolumic relaxation time (IVRT/HR) was prolonged in DOX-treated mice from the first dose (5 mg/kg) and remained elevated at 20 mg/kg, indicating early impaired active relaxation. (**D**) Medial early diastolic mitral annular velocity (e′), a relatively load-independent index of relaxation, was markedly reduced at both doses. (**E**) Transmitral filling velocities (E, A) and the E/A ratio. E and A were preserved at 5 mg/kg; the E/A ratio was unchanged at 5 mg/kg and declined at 20 mg/kg (paired 5 vs 20 mg/kg, P = 0.005), without a significant change in E or A individually. (**F**) Medial E/e′ ratio and left atrial area. E/e′ rose sharply at 5 mg/kg, driven by the fall in e′ rather than elevated filling pressures, while left atrial area remained normal at both doses, arguing against elevated filling pressures or atrial remodeling. Data are mean ± S.D (n=9-10 control, 10 DOX). Statistics: two-way repeated-measures ANOVA (treatment × dose), with Šídák-corrected multiple comparisons. *P < 0.05, **P < 0.01, ***P < 0.001, ****P < 0.0001; ns, not significant.

Heart rate and LVEF did not differ between groups at either dose (**Figure 2B**). Impaired active relaxation was the earliest abnormality, already present after the first 5 mg/kg exposure. Heart rate-corrected isovolumic relaxation time (IVRT/HR) was prolonged in DOX mice from 5 mg/kg (**Figure 2C**; *p<0.01 vs control*) and remained elevated at 20 mg/kg (*p<0.001 vs control*), without further change between doses (*paired 5 vs 20 mg/kg, p=0.97*). In parallel, medial early diastolic mitral annular velocity (e′), was markedly reduced at both doses (**Figure 2D***; p<0.0001 vs control at both doses*). Thus, two independent indices of active relaxation were concordantly impaired from a single DOX exposure (interaction/treatment; two-way repeated-measures ANOVA, p<0.05).

By contrast, transmitral flow were preserved at 5 mg/kg. Neither E nor A (**Figure 2E**) differed from control, and E/A was unchanged (**Figure 2E;** *p=0.29*). The medial E/e′ ratio, however, rose sharply at 5 mg/kg (**Figure 2F;** *p<0.0001 vs control*). As E was unchanged, this increase was driven entirely by the fall in e′ and reflected impaired relaxation rather than elevated filling pressures. Consistent with this, left atrial area was unchanged at both doses (**Figure 2F**), arguing against established elevated filling pressures or atrial remodeling.

At 20 mg/kg, the relaxation defect persisted (IVRT and e′; **Figure 2C, D**) and the transmitral filling pattern began to shift, with E/A ratio declining in DOX mice (**Figure 2E;** *paired 5 vs 20 mg/kg, p=0.005; DOX vs control at 20 mg/kg, p=0.069*), without significant change in either E or A individually (**Figure 2F**), and with no change over time in controls (*p=0.43*), consistent with the combined effect of small concordant shifts in both velocities rather. Medial E/e′ ratio returned toward control values at 20 mg/kg (**Figure 2F**), mirroring the partial rise in e′; which nonetheless remained well below control; left atrial area remained normal throughout DOX treatment (**Figure 2F**).

Together, these data delineate a temporal hierarchy of diastolic impairment. Active myocardial relaxation (IVRT, e′) is impaired from a single DOX exposure, while systolic function, transmitral filling velocities and atrial size are preserved; the filling pattern (E/A) begins to deteriorate only at the higher cumulative dose, still without atrial remodeling. This identifies impaired active relaxation as the earliest functional manifestation of anthracycline cardiotoxicity.

### Doxorubicin induces early, hierarchical spatial remodeling of cardiomyocyte subsarcolemmal and interfibrillar mitochondria

Because active myocardial relaxation is energy-dependent, we reasoned that this early defect upon DOX treatment might reflect an underlying alteration of cardiomyocyte mitochondria, and focused on the CM surface, where SSM reside within lateral crest structures and are selectively more vulnerable to cardiac stress compared to IFM^2,3^, likely owing to their location at the surface.

To resolve mitochondrial architecture at the CM surface, we used atomic force microscopy (AFM) in force-volume mode on living adult CMs, as we previously described^2,3^, simultaneously mapping surface topography and local elasticity. We previously established, in myocardial infarction (MI)-induced HF, that the CM surface displays distinct topographic/elastic signatures corresponding to specific mitochondrial architectures and defining a stereotyped sequence of surface remodeling with (**i**) the presence of periodic crest/SSM structures (Crest-like pattern) in normal CMs, (**ii**) their progressive flattening upon SSM depletion (flat pattern), and (**iii**) the accumulation of detached IFM at the surface beneath the sarcolemma, preceding CM death (heap-like pattern)^2,3^. We used these reference signatures to track surface mitochondrial remodeling during DOX exposure.

Absolute elasticity values (Young’s modulus, YM) depend on multiple factors, including the AFM setup (cantilever stiffness, calibration), and are particularly variable on soft surfaces such as the living CM surface. Because this study used different instrumentation from our previous work^2^, we first re-validated the surface elasticity signatures associated with each mitochondrial pattern, using formamide-induced sarcolemmal stress on freshly isolated living adult CMs (2-month-old male C57BL/6J). As previously reported^2^, formamide-induced sarcolemmal stress recapitulates post-MI-like alterations at the CM surface, inducing SSM disruption (flat pattern) at 250 µM and IFM disorganization with sarcolemmal displacement (heap-like pattern) at 750 µM. Consistent with these structural changes, control CMs exhibited low surface elasticity (YM ∼4 kPa), characteristic of the crest/SSM pattern, whereas 250 µM formamide reduced elasticity to <2 kPa, corresponding to the flat pattern. In contrast, 750 µM formamide markedly increased surface stiffness (>8.5 kPa), consistent with the heap-like pattern (**Supplementary Figure 1**). These distinct elasticity signatures were therefore used as reference patterns for subsequent analysis of DOX-treated CMs.

We next assessed CM surface topography/elasticity of the CMs at baseline and during DOX or saline treatment using living adult CMs isolated at defined time points: prior to injection, 3 days after the first dose (5 mg/kg), one week after a cumulative dose of 20 mg/kg, and one week after reaching 25 mg/kg (**Figure 3A and 1A**). We first controlled the stability of the crest-like pattern at baseline (60-days-old mice) and following the fourth and fifth saline injections (120 and 130-days-old mice). As indicated in **Supplemental Figure 2**, CMs exhibited stable, homogeneous surface elasticity values around 3 kPa before and after saline injections, consistent with preserved periodic crest structures along the CM surface. These conditions were therefore pooled and used as controls for subsequent analyses. In contrast, CMs from DOX-treated mice (5, 20, or 25 mg/kg) showed a progressive increase in mean surface elasticity (**Supplemental Figure 3**). At 5 and 25 mg/kg, the distribution was broad and could be resolved into two Gaussian populations, reflecting the coexistence of crest-like, flat and heap-like patterns within single AFM maps. To resolve this heterogeneity, we developed a pattern-based segmentation approach. AFM elasticity maps were segmented according to crest-like, flat, and heap-like surface patterns (**Supplemental Figure 4**), allowing calculation of pattern-specific surface coverage and mean elasticity. For each region, both the area and the mean elasticity were calculated, allowing us to determine, for every map, the surface coverage and average stiffness specific to each pattern. Using this approach, we observed a progressive loss of the crest/SSM-like pattern from the first DOX injection (5mg/kg), concomitant with the emergence of flat and heap-like patterns (**Figure 3B**). To distinguish direct CM effects from systemic influences, similar AFM analyses were performed on CMs isolated from 2-month-old male C57BL/6J mice and exposed to DOX *in vitro* for 5 min at 37°C under 5 % CO_2_ (**Figure 3C and Supplemental Figure 5**). Consistent with *in vivo* findings, DOX induced a dose-dependent loss of the crest-like patterns at the CM surface, paralleled by transient emergence of flat regions and a sustained increase in heap-like structures. These observations support a model in which the early-progressive loss of SSM at the CM surface is paralleled by the displacement of IFM toward the surface, reflecting sequential mitochondrial remodeling in response to DOX treatment.

**Figure 3.**
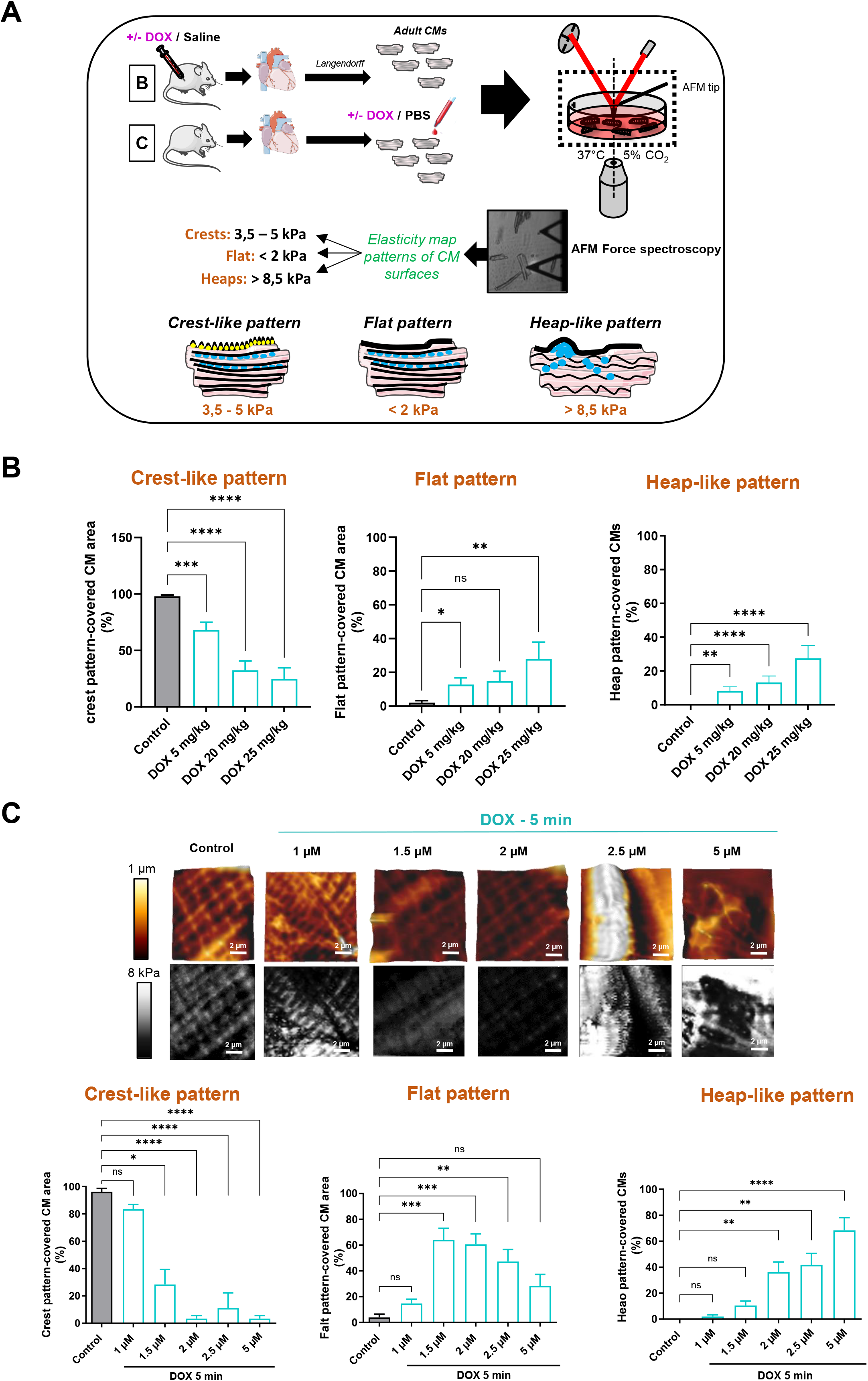
Atomic force microscopy reveals early and progressive remodeling of cardiomyocyte surface architecture and stiffness upon cumulative DOX exposure. **(A)** Schematic representation of the experimental workflow. CMs were isolated from adult male mouse hearts following either *in vivo* (**B**) or *in vitro* (**C**) doxorubicin (DOX) exposure. *In vivo*, mice received weekly intraperitoneal DOX (cumulative 5, 20 or 25 mg/kg), followed by Langendorff perfusion and CM isolation after each dose. *In vitro*, freshly isolated adult CMs CMs from 2-month-old male mice were exposed to increasing DOX concentrations (1-5 µM) for 5 min at 37°C under 5% CO₂. In both settings, surface topography and stiffness of living CMs were assessed by atomic force microscopy (AFM) in force-volume mode under physiological conditions (37°C, 5% CO₂). Three CM surface patterns were defined by their elastic signatures: crest-like (intermediate elasticity, 3.5-5 kPa), flat (soft, <2 kPa) and heap-like (stiff, >8.5 kPa). (**B**) Relative proportion of each surface pattern in CMs from mice exposed *in vivo* to increasing cumulative DOX doses or saline (control). The crest-like pattern was progressively lost from the first dose (5 mg/kg), with concomitant emergence of flat and heap-like patterns. (**C**) Relative proportion of each surface pattern in CMs exposed *in vitro* to increasing DOX concentrations or vehicle (control). *Upper panels:* representative AFM 3D topography (top) and corresponding elasticity maps (bottom) illustrating crest-like (left), flat (middle) and heap-like (right) patterns. Acute DOX reproduced the dose-dependent loss of the crest-like pattern seen *in vivo*. Data are mean ± S.E.M (*In vivo*: n=5 mice/group, ∼15 CMs/mice; *In vitro*: n=3 mice/group, 9 CMs/mice). Statistics: Kruskal-Wallis test followed by Dunn’s multiple comparisons test to compare each DOX-treated groups to the control. * *P* <0.05, ** *P* <0.01, *** *P* <0.001, **** *P* <0.0001. ns: not significant.

We confirmed this early remodeling of SSM and delayed IFM alterations during DOX-treatment by transmission electron microscopy (TEM) on left ventricular tissue across the treatment course as in **Figure 1A**. We observed a progressive reduction in crest heights at the CM surface over the course of DOX treatment, beginning at early DOX doses (20 mg/kg; **Figure 4A**), paralleled by a significant, dose-dependent decline in SSM area (SSM shrinking) (**Figure 4B**) and SSM number (**Figure 4C**). These early, progressive SSM alterations mirrored the gradual loss of mature crest/SSM at the CM surface (**Figure 4D**), consistent with the early disappearance of the crest-like pattern and the emergence of the flat pattern observed by AFM. At later stages, in agreement with the progressive emergence of heap-like surface patterns detected by AFM, TEM revealed intracellular IFM disorganization, and accumulation of IFM (heaps) beneath the sarcolemma (**Figure 4D**, left panels, DOX 25 mg/kg+15weeks). In parallel, quantification of inter-CM lateral space, reflecting crest-crest lateral interactions between adjacent CMs and as an indicator of cardiac tissue cohesion^1,3^, showed a mild increase after cumulative DOX 25 mg/kg, and a significant enlargement after 15 weeks (**Figure 4E**), coinciding with the onset of systolic dysfunction (**Figure 1B**).

**Figure 4.**
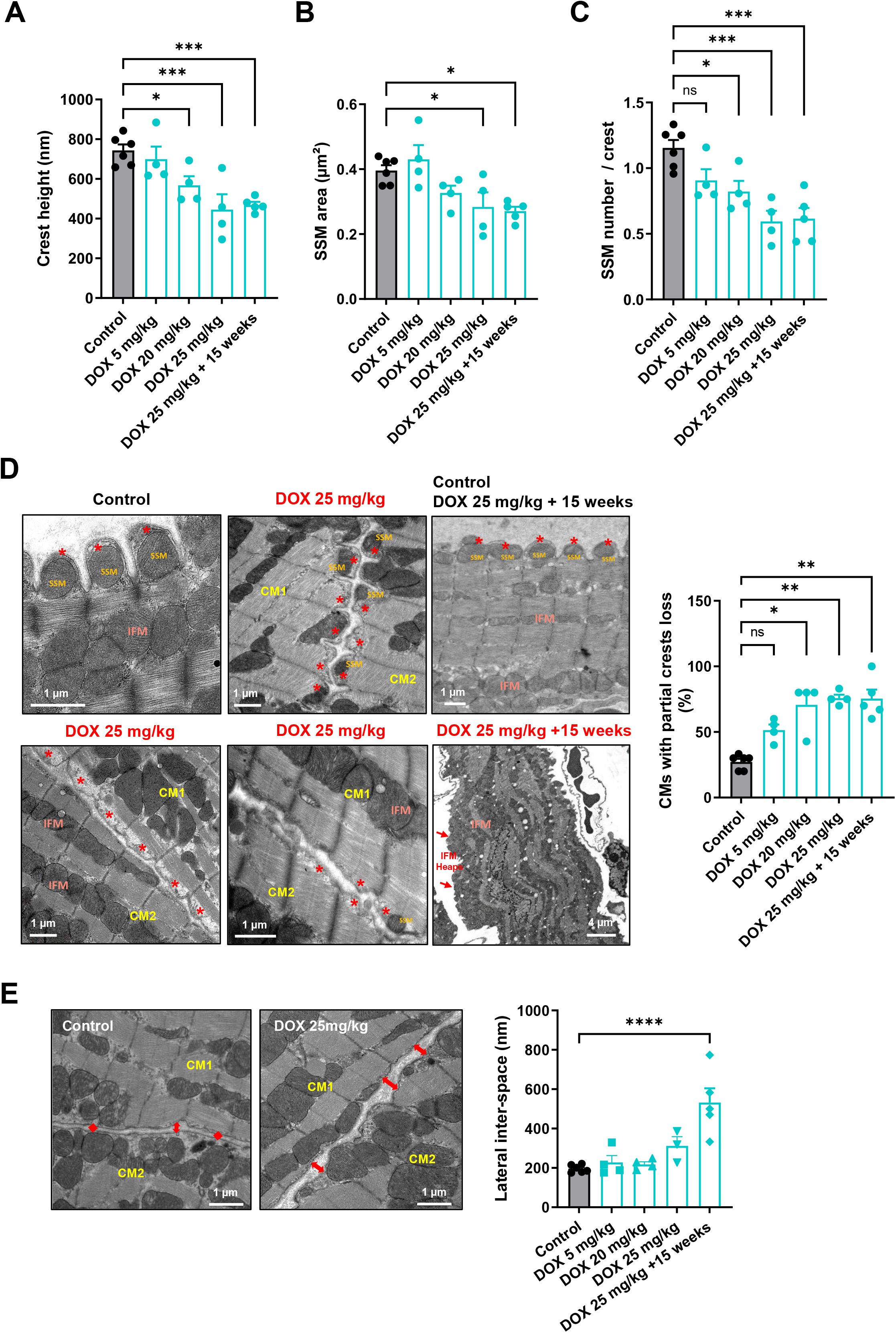
Transmission electron microscopy confirms early/progressive loss of surface crests/SSM in cardiomyocytes, preceding interfibrillar mitochondrial disorganization, upon cumulative DOX exposure. TEM-based quantification of (**A**) crest height, (**B**) subsarcolemmal mitochondrial (SSM) area, (**C**) SSM number per crest, (**D**) the proportion of CMs with partial crest loss, and (**E**) the lateral inter-CM space, in left ventricular tissue from control (saline) or DOX-treated mice (5, 20, 25 mg/kg and 25 mg/kg + 15 weeks; protocol as in Figure 1A). Crest height, SSM area and SSM number declined progressively and dose-dependently; the lateral inter-CM space widened significantly at 15 weeks. (**D**, *Left panels*) Representative TEM images showing CM surface crests and SSM in control mice (red stars), SSM loss/shrinkage in DOX 25 mg/kg-treated mice (red stars), and, at 15 weeks, intracellular interfibrillar mitochondrial (IFM) disorganization with surface IFM accumulation (IFM “heaps”; red arrows). Data are mean ± S.E.M. (n=3-4 mice/group; ≥30 crests/mouse). Statistics: one-way ANOVA followed by Dunnett’s multiple comparisons test to compare each DOX-treated groups to the control (A, B, C, E); Kruskal-Wallis test with Dunn’s multiple comparisons to compare each DOX-treated groups to the control test (D). * *P* <0.05, ** *P* <0.01, *** *P* <0.001, **** *P* <0.0001. ns: not significant.

Finally, to confirm the early structural loss of surface crest/SSM depicted by AFM et TEM during initial DOX treatment at the molecular level, we quantified the crest/SSM markers Ephrin-B1 and Claudin-5. As both are transmembrane proteins also expressed in CMs and endothelial cells, precluding whole-tissue western blot, we quantified their CM surface pool in fresh native left ventricular tissue by 3D fluorescence imaging that we recently developed (3D-NaissI method^4^), using a dedicated segmentation macro separating the CM surface (WGA⁺ lateral membrane, reflecting surface localization of SSM) from the interior and excluding the endothelial compartment (IsoB4; **Figure 5A**). Ephrin-B1 surface expression was significantly reduced in DOX-treated CMs from the first dose (5 mg/kg) and remained decreased at 20 mg/kg (**Figure 5B**). Claudin-5, which mediates crest–crest contacts between adjacent CMs, was likewise reduced at the CM surface at 5 mg/kg and 20 mg/kg (**Figure 5C**). Thus, the two molecular determinants of the surface crest/SSM compartment were depleted from a single 5 mg/kg DOX exposure, providing an independent molecular confirmation of the early surface SSM loss detected by AFM and TEM, and mirroring in time the impairment of active relaxation.

**Figure 5.**
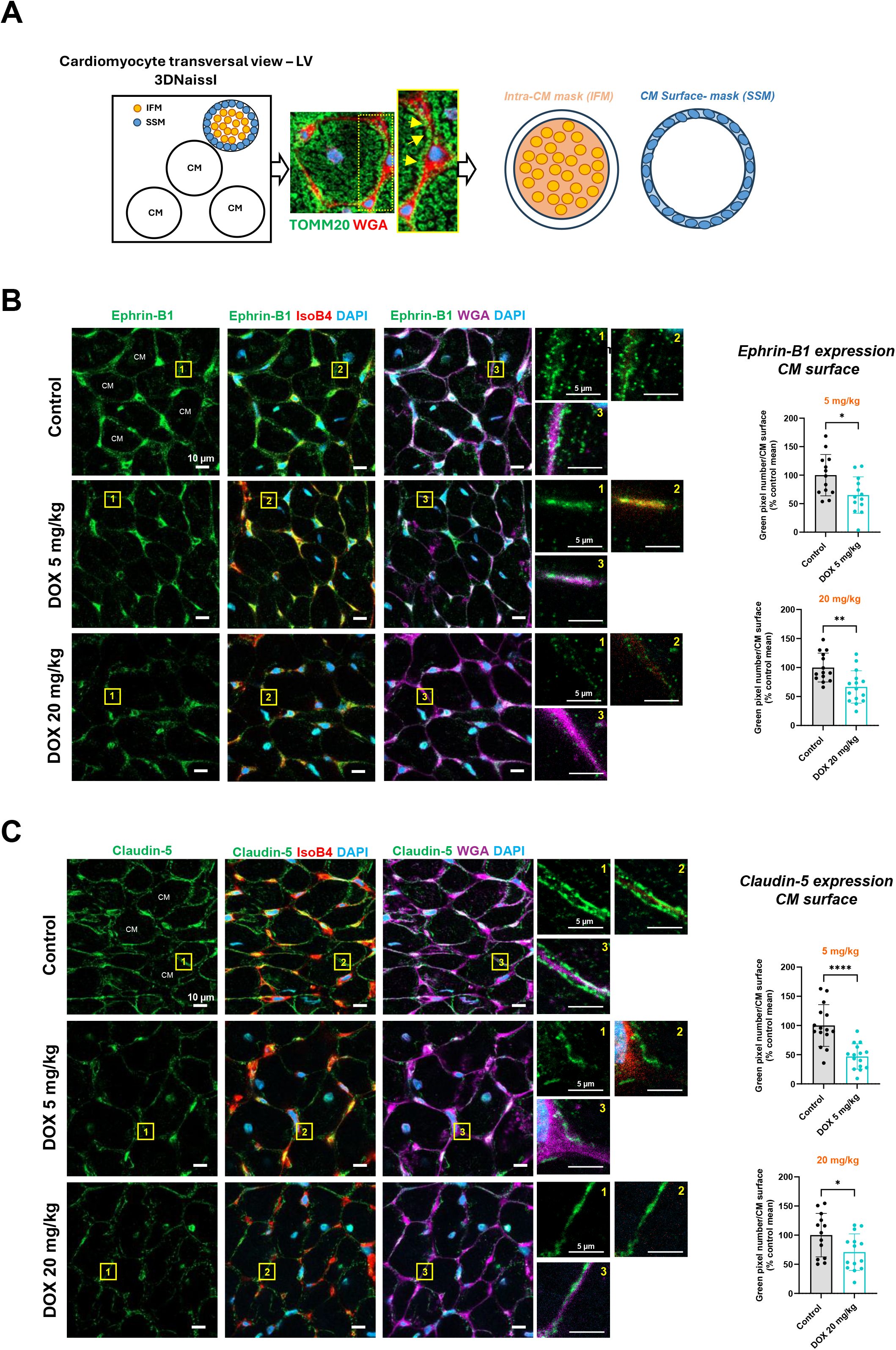
Doxorubicin depletes the crest/SSM surface determinants Ephrin-B1 and Claudin-5 from the first dose. (**A**) Principle of 3D-NaissI quantitative imaging on native left ventricular tissue (transverse cardiomyocyte/CM view). Mitochondria (TOMM20^+^, green and the lateral membrane (WGA^+^, red) were used to define two masks: a CM surface mask, capturing the WGA⁺ lateral membrane and its subsarcolemmal mitochondria (SSM; yellow arrows), and an intracellular mask, capturing interfibrillar mitochondria (IFM). The endothelial compartment (IsoB4^+^) was excluded. (**B**) Ephrin-B1 expression at the CM surface in control and DOX-treated mice. *Left:* representative fluorescent confocal images, with magnified insets of the lateral membrane. *Right:* quantification at 5 mg/kg and 20 mg/kg. Surface Ephrin-B1 was significantly reduced in DOX-treated CMs at both doses. (**C**) Claudin-5 expression at the CM surface, imaged and quantified as in (**B**). Surface Claudin-5, which mediates crest-crest contacts, was significantly reduced at both doses. Data are expressed as percentage of the control mean ± S.D. (n=3 mice; 5 biopsies/mouse; 3 stacks/biopsy; 30-75 CMs/mouse). Statistics: unpaired Student’s t-test (DOX vs control at each dose). *P < 0.05, **P < 0.01, ***P < 0.001, ****P < 0.0001; ns, not significant.

### Doxorubicin induces spatially ordered activation of PINK1/Parkin quality-control pathway, from subsarcolemmal to interfibrillar mitochondria

To determine whether the early loss of SSM and the later IFM alteration reflected a spatially ordered mitochondrial stress in CMs during DOX treatment, we examined Ser65-phosphorylated Parkin (pS65-Parkin), a marker of PINK1/Parkin-dependent quality control targeting damaged mitochondria^25,26^.

Ser65-phosphorylated Parkin (pS65-Parkin) was imaged by fluorescence within the native cardiac tissue by 3D-NaissI and quantified to separately assess the CM surface, reflecting quality control of SSM, and the CM interior, reflecting quality control of IFM (**Figure 6A, B**). CM surface (SSM) pS65-Parkin was significantly increased from the first dose (5 mg/kg) and remained elevated at 20 mg/kg (**Figure 6B**). By contrast, intra-CM IFM pS65-Parkin was unchanged at 5 mg/kg and increased only at 20 mg/kg (**Figure 6B**). Quality-control signaling was therefore activated first on SSM and only later on IFM, identifying surface SSM as the earliest site of mitochondrial stress from the first DOX dose, consistent with their early depletion detected by AFM, TEM and molecular imaging.

**Figure 6.**
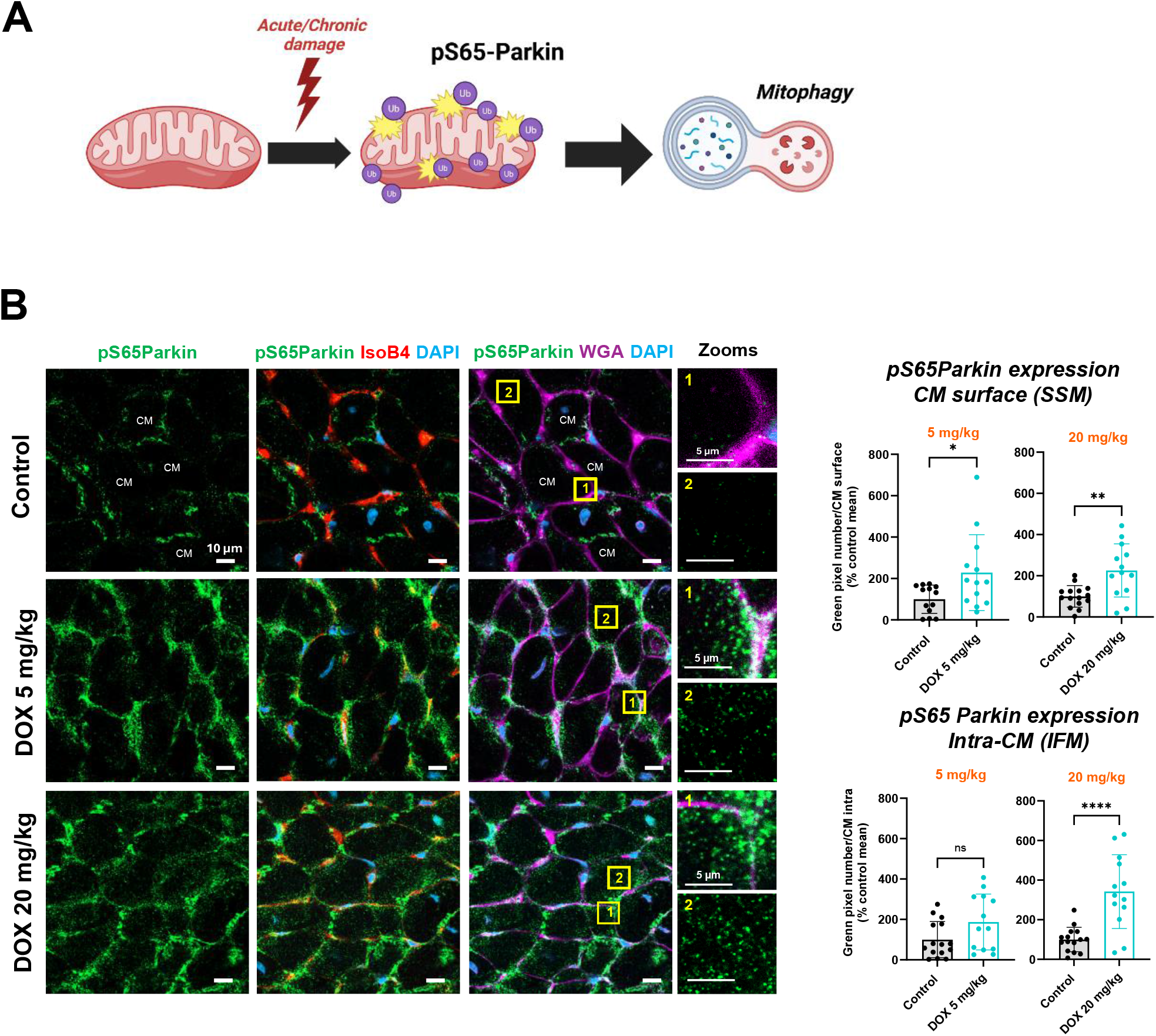
Doxorubicin induces spatially ordered activation of PINK1/Parkin quality control, engaging subsarcolemmal before interfibrillar mitochondria. (**A**) Schematic of the PINK1/Parkin mitochondrial quality-control pathway: upon mitochondrial damage, Parkin is phosphorylated at Ser65 (pS65-Parkin) and activated, promoting ubiquitination and clearance of damaged mitochondria. (**B**) pS65-Parkin expression imaged by 3D-NaissI and quantified, as explained in Figure 5A, at the CM surface (subsarcolemmal mitochondria, SSM) and CM interior (interfibrillar mitochondria, IFM), in control and DOX-treated mice. *Left:* representative fluorescent confocal images, with magnified surface and intracellular insets. *Right:* quantification at 5 and 20 mg/kg. Surface (SSM) pS65-Parkin was significantly increased from the first dose (5 mg/kg) and remained elevated at 20 mg/kg. In contrast, interior (IFM) pS65-Parkin was unchanged at 5 mg/kg and increased only at 20 mg/kg, indicating quality-control activation on SSM before IFM. Data are expressed as percentage of the control mean ± S.D. (n=3 mice; 5 biopsies/mouse; 3 stacks/biopsy; 30-60 CMs/mouse). Statistics: unpaired Student’s t-test (DOX vs control at each dose and compartment). *P < 0.05, **P < 0.01, ***P < 0.001, ****P < 0.0001; ns, not significant.

### Doxorubicin depletes sarcolemmal Na^+^/Ca^2+^ exchanger NCX1, linking cardiomyocyte surface SSM injury to impaired active relaxation

Crest/SSM loss at the CM surface and impaired active relaxation both emerged from the first DOX dose. To test whether they were mechanistically linked, we spatially imaged and quantified the expression of two diastolic Ca^2+-^handling determinants by 3D-NaissI in the native cardiac tissue, the sarcolemmal Na^+^/Ca^2+^ exchanger NCX1 which extrudes calcium at the sarcolemma during diastole and the intracellular sarcoplasmic reticulum pump SERCA2 (**Figure 7A**).

**Figure 7.**
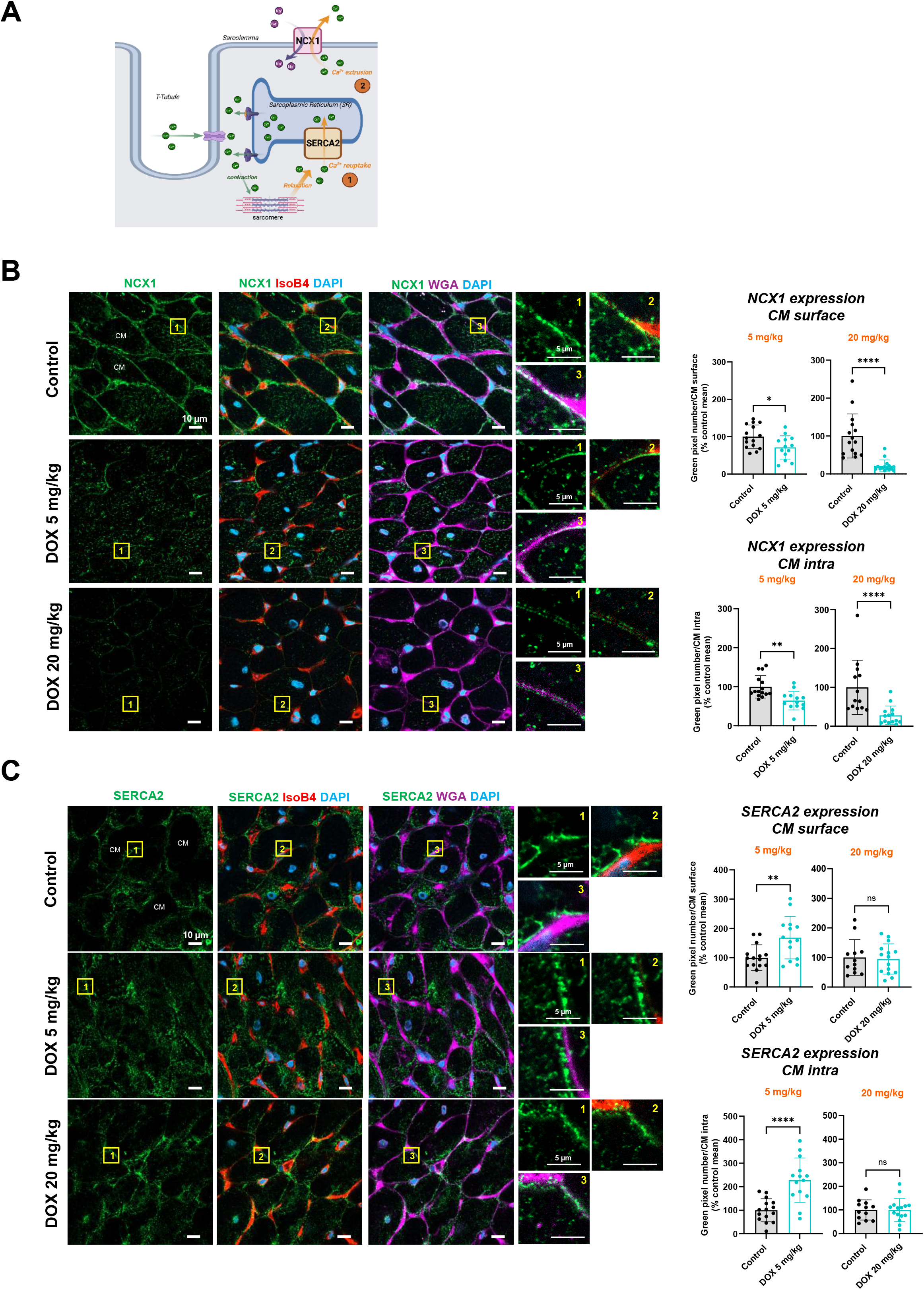
Doxorubicin depletes the sarcolemmal calcium exchanger NCX1 from the first dose, with only a transient rise in SERCA2. (**A**) Schematic of cardiomyocyte Ca^2+^ handling: during diastole, cytosolic Ca^2+^ is removed predominantly by SR reuptake through SERCA2 and, at the sarcolemma, by extrusion through the Na⁺/Ca²⁺ exchanger NCX1. (**B**) NCX1 expression quantified, as explained in Figure 5A, by 3D-NAISSI at the CM surface and interior in control and DOX-treated mice. *Left:* representative fluorescent confocal images, with magnified surface and intracellular insets. *Right:* quantification at 5 and 20 mg/kg. Surface NCX1 was significantly reduced from the first dose (5 mg/kg) and collapsed at 20 mg/kg, with a comparable decline in the CM interior. (**C**) SERCA2 expression, imaged and quantified as in (B). SERCA2 was transiently increased at both the CM surface and interior at 5 mg/kg, then returned to control levels at 20 mg/kg. Data are expressed as percentage of the control mean ± S.D. (n=3 mice; 5 biopsies/mouse; 3stacks/biopsy; 20-60 CMs/mouse). Statistics: unpaired Student’s t-test (DOX vs control at each dose and compartment). *P < 0.05, **P < 0.01, ***P < 0.001, ****P < 0.0001; ns, not significant.

CM surface NCX1 was reduced in DOX-treated CMs from the first dose (5 mg/kg) and collapsed at 20 mg/kg with a comparable decline in the CM interior (**Figure 7B**). In contrast, SERCA2 was transiently increased at both the CM surface and interior at 5 mg/kg, then returned to control levels at 20 mg/kg (***Figure 7C***). Thus, while NCX1, the sarcolemmal route of diastolic Ca^2+^ extrusion, progressively collapsed, the early SERCA2 up-regulation was self-limited and did not persist. Thus, from the first dose, the loss of sarcolemmal NCX1, only transiently offset by the rise in SERCA2, may likely impair diastolic Ca^2+^ removal and contribute to the early relaxation defect (IVRT, e′; **Figure 2**).

### Disruption of crest/SSM architecture at the cardiomyocyte surface accelerates the transition to HFrEF

Finally, we examined whether early loss of crest/SSM structures at the CM surface, associated with diastolic dysfunction, contributes to the subsequent development of systolic dysfunction following DOX exposure.

To address this question, DOX treatment was reproduced in an adult mouse model lacking mature crests/SSM at the CM surface, using a tamoxifen-inducible-CM specific-conditional-knock-out (αMHC-Mer-Cre-Mer) of *Efnb1*. We previously demonstrated that Ephrin-B1 regulates both crest/SSM and diastolic function maturation, and that its CM-specific deletion induces a HFpEF-like phenotype that progressively evolves toward HFrEF with age, with 100 % mortality observed by 13 months of age^1^. Adult male *Efnb1*^flox/flox^-αMHC-Cre^+^ mice were first treated with tamoxifen at 2 months of age to induce CM-specific *Efnb1* deletion, and one month later, allowing sufficient time to achieve approximately 50% deletion efficiency as previously reported^1^, mice were administered DOX or saline (control) at 5 mg/kg once weekly for four consecutive weeks (total cumulative dose 20 mg/kg). Cardiac function was assessed by echocardiography in each mouse longitudinally at baseline (prior to tamoxifen injection), one month after tamoxifen treatment (Tamox effect), and following DOX administration at both 5 mg/kg and 20 mg/kg time points (**Figure 8A**). Consistent with our previous findings, tamoxifen alone did not significantly alter LVEF in control mice (**Figure 8B, left panel**). In contrast, *Efnb1*-deficient mice (tamox treated) exposed to DOX exhibited a significant reduction in LVEF as early as after the first DOX injection (5 mg/kg), with further deterioration after cumulative dosing (20 mg/kg) (**Figure 8B, right panel**). Thus, whereas DOX treatment alone induces systolic dysfunction only at late stages (15 weeks after DOX treatment), pre-existing disruption of crest/SSM architecture at the CM surface markedly sensitizes the adult heart to DOX-induced injury, precipitating an accelerated transition to HFrEF.

**Figure 8.**
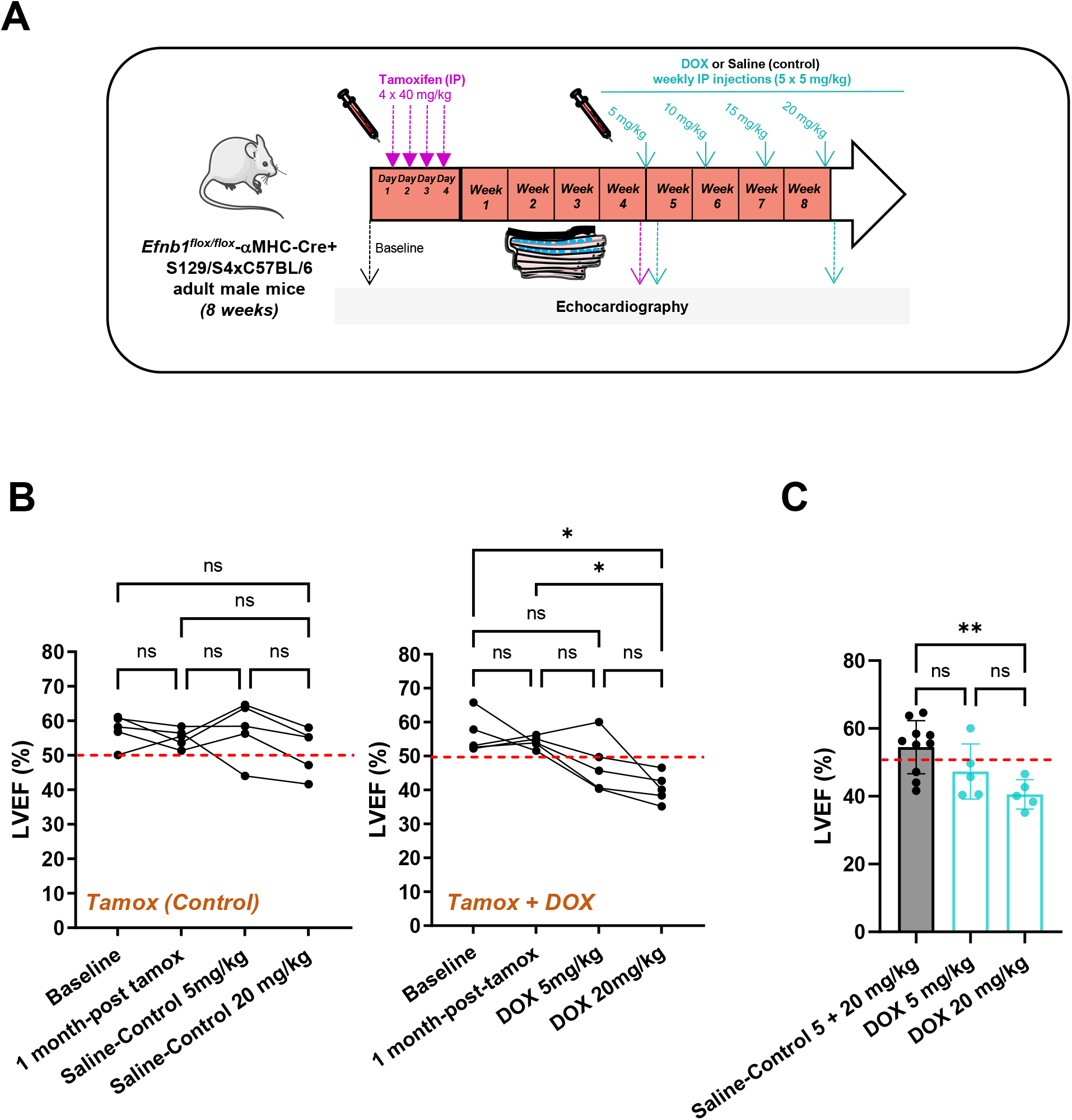
Disruption of crest/SSM structures at the cardiomyocyte surface accelerates the transition to systolic dysfunction following DOX exposure. (**A**) Schematic of the experimental protocol. Adult male *Efnb1*^flox/flox^-αMHC-MerCre^+^Mer mice (8 weeks old) received tamoxifen (4 × 40 mg/kg, i.p.) to induce CM-specific deletion of Ephrin-B1, a key regulator of surface crest/SSM maturation. One month later, allowing partial (∼50%) deletion, mice received weekly intraperitoneal DOX (5 mg/kg) or saline (control) for 4 consecutive weeks (cumulative 20 mg/kg). Transthoracic echocardiography was performed longitudinally in the same animals at baseline (before tamoxifen), one-month post-tamoxifen (Tamox effect), and after DOX or saline administration (5 mg/kg and 20 mg/kg cumulative). (**B**) Longitudinal left ventricular ejection fraction (LVEF) in Ephrin-B1-deficient mice treated with saline (*leftl*) or DOX (*right*) (n=5 mice/group). In the saline group, LVEF remained preserved after tamoxifen-induced Ephrin-B1 deletion, confirming that partial deletion alone does not impair systolic function^1^. In contrast, DOX-treated mice showed an early trend toward reduced LVEF after the first dose (5 mg/kg), worsening at 20 mg/kg, indicating that Ephrin-B1 loss sensitizes the heart to DOX-induced systolic dysfunction. (**C**) LVEF in pooled saline controls (5 + 20 mg/kg) and DOX-treated groups (5 mg/kg or 20 mg/kg), from the data in (B). Data are mean ± SD. Statistics: repeated-measures one-way ANOVA for within-mouse changes over time (**B**) and ordinary one-way ANOVA for between-group comparisons (**C**), each with Tukey’s multiple comparisons test. * P <0.05, ** P <0.01; ns: not significant.

Together, these findings identify crest/SSM as a cardioprotective mitochondrial subpopulation during the early phases of anthracycline-induced cardiotoxicity, likely buffering initial mitochondrial stress and preserving IFM organization and contractile function until their architectural displacement to the CM surface precipitates systolic failure.

## Discussion

In this study, using longitudinal, multimodal assessment from the first drug exposure, we show that a single doxorubicin (DOX) dose is sufficient to impair active myocardial relaxation, before any systolic dysfunction. This early defect selectively affected the active, energy-and Ca²⁺-dependent phase of relaxation, reflected by prolonged IVRT and reduced e′, while the passive determinants of ventricular compliance were spared. This early alteration of active relaxation is underpinned by selective injury of the surface subsarcolemmal mitochondrial (SSM) compartment of the CM, consistent across structural (AFM, TEM), molecular (Ephrin-B1, Claudin-5) and mitochondrial stress-response (PINK1/Parkin activation) readouts, and accompanied by early loss of the sarcolemmal Na⁺/Ca²⁺ exchanger NCX1, both SSM and NCX1 being effectors of active relaxation. In contrast, interfibrillar mitochondria (IFM), which mainly fuel myofibrillar contraction, were affected only later, upon cumulative DOX exposure, likely supporting later systolic dysfunction. In CM-specific *Efnb1*-deficient mice lacking mature SSM, DOX precipitated an accelerated transition to systolic failure, identifying surface SSM as an early cardioprotective compartment. Together, these findings define impaired active relaxation as the earliest hallmark of anthracycline cardiotoxicity and define its subcellular origin (**Figure 9**).

**Figure 9.**
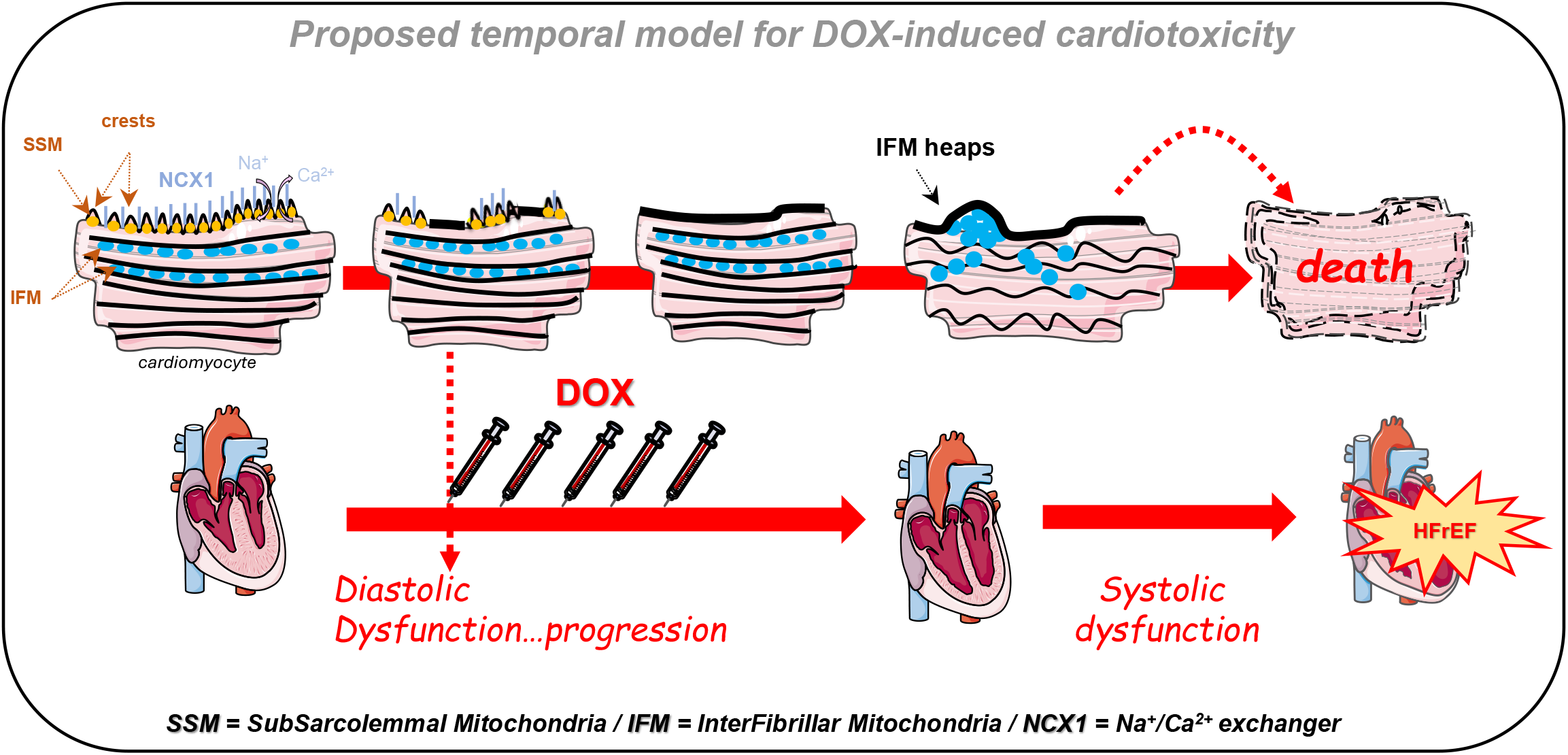
Proposed temporal model of cardiomyocyte surface remodeling in doxorubicin-induced cardiotoxicity. Schematic of the sequential alterations in cardiomyocyte (CM) mitochondrial architecture underlying the progression from early diastolic dysfunction to systolic failure during DOX cardiotoxicity. Under physiological conditions, subsarcolemmal mitochondria (SSM) reside within periodic surface crest structures, alongside the sarcolemmal Na⁺/Ca²⁺ exchanger NCX1, supporting local bioenergetics, diastolic calcium extrusion and lateral tissue cohesion. From the first DOX dose, this surface crest/SSM microdomain is selectively and progressively lost, together with sarcolemmal NCX1, impairing active relaxation while ejection fraction, transmitral filling and atrial size remain preserved, an early diastolic phenotype without established filling-pressure elevation or structural remodeling. This surface injury is accompanied by spatially ordered activation of the PINK1/Parkin mitochondria quality-control pathway, engaged on surface SSM before deeper interfibrillar mitochondria (IFM) (Figure 6), consistent with early targeting of CM surface mitochondria. Upon sustained injury, IFM, which primarily energize myofibrillar contraction, in turn detach from the contractile apparatus and aggregate into heap-like structures beneath the sarcolemma. This disorganization coincides with mitochondrial failure and CM death, as previously reported², driving progression toward systolic dysfunction and heart failure with reduced ejection fraction (HFrEF). Overall, this model positions early remodeling of the CM surface crest/SSM/NCX1 microdomain, and the associated impairment of active relaxation, as the initiating event of DOX cardiotoxicity.

Early diastolic abnormalities have been reported clinically during anthracycline therapy, but over repeated cycles and without a defined cellular substrate. Our data extend this concept by showing that i/ impaired active relaxation is detectable from the very first DOX dose and ii/ it selectively involves the active phase of relaxation, with an identifiable subcellular origin in the CM surface SSM compartment and its sarcolemmal Ca²⁺ effector NCX1 Collectively, these results support systematic diastolic monitoring during anthracycline therapy as a sensitive strategy for early detection and timely cardioprotection

### Diastolic dysfunction as a primary feature of DIC

Although reduced LVEF has historically defined DIC, a multiparametric approach is increasingly used to detect early injury before a detectable decline in LVEF, including GLS^27–29^, circulating cardiac troponins^12,30,31^, and cardiac magnetic resonance^32–34^. Our study shows that impaired active relaxation, particularly IVRT prolongation and reduced e′, is detectable from a single DOX dose, in parallel with GLS abnormalities and while LVEF, transmitral filling and atrial size remain preserved. Importantly, this early phenotype reflects impaired active relaxation rather than a passive, fibrotic HFpEF-like process since E/A and left atrial size were preserved and fibrosis was minimal at this stage, arguing against established filling-pressure elevation or structural remodeling. These findings are consistent with emerging clinical observations suggesting that impaired myocardial relaxation may represent one of the earliest functional manifestations of DOX cardiotoxicity^17,35,36^, a concept that remains to be firmly established. They support the early incorporation of Doppler-based diastolic assessment into cardio-oncology follow-up, particularly in patients with preserved LVEF but subclinical dysfunction evidenced by GLS or troponin. This is especially relevant as many anthracycline-treated patients are older with comorbidities predisposing to HFpEF, in whom DOX injury may exacerbate an underlying diastolic vulnerability.

### Subsarcolemmal mitochondria as the earliest subcellular target of DIC

While reduced GLS^27–29^ and elevated circulating troponin^12,30,31^ are recognized as early indicators of anthracycline cardiotoxicity, their cellular basis has remained undefined. We show that early DOX-induced relaxation impairment is tightly associated with spatially restricted remodeling of the surface crest/SSM compartment, evident from the first dose across three independent approaches including structural (crest flattening by AFM, SSM shrinkage and loss by TEM), molecular (depletion of the crest/SSM determinants Ephrin-B1 and Claudin-5), and stress-response (CM surface PINK1/Parkin activation). Notably, the early loss of Claudin-5 parallels the progressive enlargement of the inter-CM lateral space, consistent with a loss of crest-crest cohesion at the lateral CM surface. These results extend previous reports identifying metabolic disturbances as early features of DIC^37–39^ and provide structural correlates that mechanistically connect mitochondrial injury to functional impairment. This early SSM loss coincides with CM atrophy, in line with our prior work showing that physiological crest/SSM maturation through SSM swelling supports an atypical hypertrophic program independent of myofibrillar growth^1^; its rapid regression may thus represent an early, potentially reversible response to subclinical stress. Importantly, we provide direct evidence that SSM are affected earlier than IFM, whose structural integrity remains preserved at stages when diastolic dysfunction is already evident. Preferential SSM vulnerability has already been reported in myocardial ischemia^2^ and pressure overload-induced hypertrophy^3^, reinforcing the concept that mitochondrial subpopulations display distinct stress sensitivities.

### Spatial and temporal hierarchy of cardiomyocyte mitochondrial injury in DIC

This study reinforces the underappreciated concept that SSM and IFM within CMs not only serve distinct physiological roles but also exhibit temporally distinct stress vulnerabilities. The early disruption of SSM, localized just beneath the sarcolemma, contrasts with the delayed remodeling of deeper IFM, a sequence directly demonstrated by the spatially ordered activation of the PINK1/Parkin quality-control pathway, engaged first on SSM (from the first dose) and only later on IFM. This hierarchy mirrors, in reverse, their physiological developmental trajectories. Hence, IFM mature early after birth^40^ and primarily support systolic contraction through ATP production and calcium buffering near the myofibrils whereas SSM undergo delayed postnatal maturation at the CM sarcolemma and contribute to diastolic function and localized energetic support^1^. The early and selective depletion of SSM at the CM surface likely compromises first the energetic and effector machinery required for efficient relaxation, specifically sarcolemmal NCX1, before stress propagates to the broader mitochondrial network and contractile function. This spatial compartmentalization of CM mitochondria cannot be captured by bulk biochemical approaches or by studies of isolated SSM and IFM given the lack of specific markers, and implies that preserving SSM integrity early could delay downstream IFM remodeling and systolic failure, a potential therapeutic window.

### Sarcolemmal NCX1 loss links surface SSM injury to impaired active relaxation

Efficient diastolic relaxation requires rapid removal of cytosolic Ca^2+^, which is mediated predominantly by SR reuptake through SERCA2 and, to a lesser extent, by sarcolemmal extrusion through the Na⁺/Ca²⁺ exchanger NCX1^41^. In overt HF, SERCA2 activity is reduced whereas NCX1 is generally upregulated as a compensatory extrusion pathway, although the contribution of NCX1 to diastolic dysfunction remains debated^42^. Here, we observed here the opposite pattern. Surface NCX1 was reduced from the first DOX dose and further decreased at higher cumulative exposure, whereas SERCA2 showed only a transient increase that returned to control levels. This early loss of NCX1 is therefore unlikely to represent the compensatory remodeling of established HF, and more probably results from the structural disruption of the sarcolemmal microdomain in which NCX1 is localized, namely the crest/SSM compartment that was selectively lost in our model. Although Ca^2+^ handling was not directly assessed in this study, the early reduction of this sarcolemmal extrusion pathway, incompletely compensated by the transient SERCA2 increase, provides a plausible substrate for impaired diastolic Ca^2+^ removal and for the concomitant early relaxation defect (IVRT, e′; **Figure 2**).

More broadly, the diastolic determinants altered in our DOX model, *i.e.* sarcolemmal mitochondrial energetics (SSM), sarcolemmal Ca²⁺ extrusion (NCX1) and SR reuptake (SERCA2), all belong to the energy- and Ca²⁺-dependent machinery of active relaxation. In contrast, the passive determinants of ventricular compliance, such as interstitial collagen and titin-dependent myocardial stiffness, were not affected, as suggested by the preserved E/A ratio and left atrial size and by the minimal fibrosis observed at this stage. Early anthracycline cardiotoxicity thus appears to affect primarily the active phase of diastole, consistent with the functional phenotype (prolonged IVRT and reduced e′) and distinct from a passive, fibrotic HFpEF-like process.

### Proposed model for the temporal evolution from diastolic to systolic dysfunction in DIC

Our findings support a model in which early disruption of the CM surface crest microdomain, integrating SSM, junctional (Claudin-5, Ephrin-B1) and Ca^2+-^handling (NCX1) determinants, initiates diastolic dysfunction, while subsequent intracellular remodeling drives progression to systolic failure. We have previously shown that lateral membrane structure and biomechanics of the CM are regulated by Ephrin-B1 in complex with Claudin-5^24^, a pathway critical for crest maturation, inter-CM cohesion, and diastolic function^1^. The early, quantified loss of both proteins under DOX indicates disruption of this axis. Thus, Claudin-5 loss weakens crest–crest cohesion, whereas Ephrin-B1 loss likely compromises lateral membrane resilience and may impair SSM energetic function through mechanisms that remain to be elucidated. Over time, repetitive contraction on this fragile substrate may propagate stress to intracellular T-tubules and IFM, culminating in IFM detachment, CM death and systolic dysfunction. Consistent with this model, DOX in CM-specific *Efnb1*-deficient mice, with immature SSM, reduced cohesion and a HFpEF-like phenotype^1^, precipitates rapid transition to HFrEF. Supporting translational relevance, *Efnb1* is downregulated in patients with HFpEF and HFrEF, more markedly in HFpEF⁴¹, and correlates with systolic function in experimental pressure overload^24^.

### Study limitations

This study assessed DOX cardiotoxicity in a murine model without an underlying cancer or tumor-related microenvironment, and therefore does not fully replicate the clinical setting of oncology patients. In addition, only male mice were studied, limiting extrapolation to patient heterogeneity (sex differences, comorbidities, concomitant anticancer therapies). Moreover, we characterized the structural, molecular and spatial organization of mitochondrial subpopulations rather than their bioenergetic function, as direct functional assays of isolated SSM and IFM remain technically challenging. Up to date, no reliable specific markers discriminate between these two mitochondria subpopulations, underscoring the value of the spatially resolved imaging used here. Likewise, we quantified NCX1 and SERCA2 expression rather than Ca^2+^ handling itself, and Parkin activation rather than mitophagic flux.

### Perspectives

While GLS is an established marker of subclinical cardiotoxicity, our findings support that diastolic function provides additional and complementary information for early detection, from the first anthracycline treatment. Recognizing both parameters may improve risk stratification and timely cardioprotective interventions in patients receiving anthracyclines. Future clinical studies should determine whether integrating diastolic assessment into GLS-based monitoring improves early detection and outcomes, and whether strategies preserving surface SSM integrity can delay or prevent progression to HF in patients receiving anthracyclines.

## Acknowledgements

We are grateful to TRI Genotoul network facilities in Toulouse, particularly the ANEXPLO platform (Toulouse) for assistance with echocardiography, and the “Centre de Microscopie Electronique Appliquée à la biologie-CMEAB» (Faculté de Médecine Rangueil, Toulouse). We also thank Marine Josse for her contribution to TEM analysis. We acknowledge the « I2MC - Plateau Histologie Imagerie » (Genotoul-TRI), member of the national infrastructure France-BioImaging for confocal imaging of fresh cardiac biopsies.

## Author Contributions

C.G.F. and C.K. designed experiments, performed and analyzed cardiac function and transmission electron microscopy (TEM) data, and supervised the overall project. R.G. performed, analyzed and interpreted the diastolic functional analyses, and contributed to writing the manuscript. M.C. performed and analyzed *in vivo* doxorubicin (DOXO) and TEM experiments, and contributed to cardiac functional analysis. V.L. performed and analyzed atomic force microscopy (AFM) experiments and contributed to data interpretation. N.P. contributed to the *in vivo* DOX experiments and carried out the 3D-NAISSI imaging/analysis of fresh cardiac biopsies. O.L. performed, analyzed, and interpreted part of the cardiac function experiments. E.D. designed, supervised, and interpreted the AFM experiments. C.S. analyzed and interpreted the AFM datasets. J.M.S. conceived the overall DOXO protocol in mice to mimic the clinical setting, contributed to data interpretation, and critically revised the manuscript. C.G. conceived and designed the study, analyzed data, integrated all experimental findings, and wrote the manuscript with input from all authors.

The authors used BioRender.com to create Schematic illustrations in the Figures and Anthropic Claude for English language editing only.

## ABBREVIATIONS AND ACRONYMS

AFM: Atomic force microscopy
CM: Cardiomyocyte
DIC: Doxorubicin-induced cardiotoxicity
HF: Heart Failure
HFrEF: Heart Failure with reduced ejection fraction
HFpEF: Heart Failure with preserved ejection fraction
IFM: Interfibrillar mitochondria
IVRT: Isovolumic relaxation time
cKO: Conditional Knock-out
LVPWd: Left ventricular posterior wall in diastole
LVEDV: Left ventricular end-diastolic volume
LVEDD: Left ventricular end-diastolic diameter
LVIDd: Left ventricular internal diameter in diastole
LV-GLS: Left ventricular global longitudinal strain
SSM: Subsarcolemmal mitochondria
TEM: Transmission electron microscopy
WT: Wild-type

## SUPPLEMENTAL FIGURE LEGENDS

**Supplemental Figure 1. Definition of cardiomyocyte (CM) surface patterns by their elasticity signatures.**

(**A**) Schematic representation of the experimental protocol. Freshly isolated cardiomyocytes (CMs) from adult male C57BL/6J mice (8 weeks) were obtained by Langendorff perfusion and analyzed by atomic force microscopy (AFM) in force-volume mode (37°C, 5% CO₂). To reproduce the three CM surface patterns previously identified *in vivo* and validate their detection on a new AFM setup, isolated CMs were exposed for 20 min to increasing formamide concentrations, as previously described^2^. Each concentration was selected for its ability to reliably induce a specific pattern: no formamide (crest-like), 250 µM (flat) and 750 µM (heap-like).

(**B**) Representative elasticity maps (64 × 64 force curves over a 100 µm² area) illustrate the three main CM surface patterns and their corresponding Young’s modulus (YM): Crest-like pattern (control, untreated CMs): organized surface topography and intermediate stiffness (YM = 4.05 ± 1.8 kPa; range: 3.5-5 kPa); Flat pattern (Formamide 250 µM): reduced topographical complexity and decreased stiffness (YM = 1.3 ± 0.5 kPa; YM < 2 kPa); Heap-like pattern (Formamide 750 µM): prominent surface protrusions and increased stiffness (YM = 10.7 ± 2.2 kPa; YM > 8.5 kPa). n = 1 mouse, 3 CMs, 12,288 force curves per condition. Scale bars, 2 µm. Data are mean ± SD.

**Supplemental Figure 2. Cardiomyocyte surface architecture and stiffness are stable over time under control conditions.**

Freshly isolated cardiomyocytes (CMs) from saline-injected adult male C57BL/6J mice (8 weeks old) were analyzed by atomic force microscopy (AFM) in force-volume mode at successive time points (60, 120 and 130 days).

(**Top**) Representative AFM topography images (10 × 10 µm scan area) showing preserved surface morphology across time points

(**Middle**) Corresponding elasticity maps (4,096 force curves/cell; 10×10 µm) showing a homogeneous elasticity distribution.

(**Bottom**) Frequency distribution of Young’s modulus (YM) from all force curves, with Gaussian fitting to determine the peak YM/group (red). Surface stiffness remained stable across time points (YM≈3-4 kPa; crest-like range). n= 61,440 curves/time point; 12-15 CMs from 4-5 mice/condition. Data are mean ± SD.

**Supplemental Figure 3. Qualitative and quantitative assessment of cardiomyocyte surface stiffness remodeling following in vivo doxorubicin treatment.**

Freshly isolated adult CMs from mice exposed to increasing cumulative DOX doses (5 to 25 mg/kg) were analyzed using atomic force microscopy (AFM) in Force Spectroscopy mode. For each condition, three levels of analysis are shown: (*top*) Representative surface topography (10 µm × 10 µm); (*middle*) Elasticity map (10 µm × 10 µm; 4096 force curves); (*bottom*) Frequency distribution of Young’s modulus (YM) extracted from individual force curves. Gaussian models were applied to each distribution to extract peak YM values.

Control CMs (saline-treated, combined day 60-120-130) exhibit a unimodal distribution centered at 3.65 ± 1.8 kPa (n=50 cells, 18 mice, 209,800 curves). Following low-dose DOX exposure (5 mg/kg, day 60), the YM distribution shifted to a bimodal profile, with distinct peaks at 1.46 ± 0.7 kPa and 4.19 ± 2.6 kPa (n=21 cells, 5 mice, 86,016 curves), indicative of surface softening in a subset of CMs. At a higher cumulative dose (25 mg/kg, day 130), YM values increased, yielding a broader unimodal distribution centered at 4.9 ± 2.1 kPa (n=20 cells, 5 mice, 81,920 curves), suggesting progressive stiffening consistent with IFM aggregation at the cell surface.

**Supplemental Figure 4. Strategy for analyzing heterogeneous cardiomyocyte surface elasticity maps using motif-specific segmentation.**

(**Left**) Representative elasticity map (10 × 10 μm²) from a CM isolated 36 days after a cumulative DOX dose of 30 mg/kg, showing a bimodal Young’s modulus (YM) distribution overlaid on the image (green line). Each peak in the YM distribution corresponds to a distinct surface motif previously defined in Figure S1. Based on color-coded elasticity ranges, regions matching crest-like and heap-like motifs were segmented and analyzed separately.

(**Right**) Schematic of the segmentation approach applied to each elasticity map, assigning pixels to one of three predefined motifs (crest, flat, or heap) based on YM thresholds and topographical features. For each motif, mean elasticity (kPa) and corresponding surface area (µm²) were computed and recorded in a structured results table (excerpt shown at bottom). In the example shown, the analyzed map revealed coexisting crest and heap motifs with average elasticities of 4.12 ± 0.76 kPa and 10.8 ± 1.2 kPa, and corresponding surface areas of 57.56 µm² and 39.58 µm², respectively. This motif-specific analysis was systematically applied across all conditions to quantify spatially heterogeneous mechanical remodeling of the CM surface during DIC progression.

**Supplemental Figure 5. I*n vitro* exposure to doxorubicin induces dose-dependent alterations in cardiomyocyte surface stiffness.**

Freshly isolated CMs from adult male C57BL/6J mice (8 weeks old) CMs were incubated for 5 minutes at 37°C with increasing concentrations of DOX (1-5 µM). Cell surface mechanical properties were assessed using AFM in Force Spectroscopy mode over a 10 × 10 µm² area (64 × 64 force curves; total of 36,864 curves/condition; n = 9-19 cells; 3 mice/group).

(**Top**) Representative 3D surface topography images for each condition.

(**Middle**) Corresponding elasticity maps showing spatial distribution of Young’s modulus (YM). (**Bottom**) Frequency histograms of YM values extracted from AFM force curves. Gaussian curve fitting was used to determine the predominant YM values. Control CMs (untreated) exhibited a unimodal distribution centered at 3.6 ± 0.5 kPa; At 1.5 µM DOX, YM increased to 4.2 ± 2.6 kPa; At 2 µM DOX, YM shifted toward lower stiffness (1.5 ± 0.6 kPa); At 2.5 µM DOX, YM increased to 4.7 ± 2.2 kPa; At 5 µM DOX, the distribution became bimodal, with peaks at 3.7 ± 0.8 kPa and 7.2 ± 1.3 kPa, indicating the emergence of localized stiffened subdomains on the CM surface.

## References

1. Karsenty C, Guilbeau-Frugier C, Genet G, Seguelas MH, Alzieu P, Cazorla O, et al. Ephrin-B1 regulates the adult diastolic function through a late postnatal maturation of cardiomyocyte surface crests. Elife 2023;12. doi: 10.7554/eLife.80904

2. Dague E, Genet G, Lachaize V, Guilbeau-Frugier C, Fauconnier J, Mias C, et al. Atomic force and electron microscopic-based study of sarcolemmal surface of living cardiomyocytes unveils unexpected mitochondrial shift in heart failure. J Mol Cell Cardiol 2014;74:162–172. doi: 10.1016/j.yjmcc.2014.05.006

3. Guilbeau-Frugier C, Cauquil M, Karsenty C, Lairez O, Dambrin C, Payre B, et al. Structural evidence for a new elaborate 3D-organization of the cardiomyocyte lateral membrane in adult mammalian cardiac tissues. Cardiovasc Res 2019;115:1078–1091. doi: 10.1093/cvr/cvy256

4. Pataluch N, Guilbeau-Frugier C, Pons V, Wahart A, Karsenty C, Senard JM, et al. Unveiling the native architecture of adult cardiac tissue using the 3D-NaissI method. Cell Mol Life Sci 2025;82:70. doi: 10.1007/s00018-025-05595-y

5. Lyon AR, Lopez-Fernandez T, Couch LS, Asteggiano R, Aznar MC, Bergler-Klein J, et al. 2022 ESC Guidelines on cardio-oncology developed in collaboration with the European Hematology Association (EHA), the European Society for Therapeutic Radiology and Oncology (ESTRO) and the International Cardio-Oncology Society (IC-OS). Eur Heart J 2022;**43**:4229–4361. doi: 10.1093/eurheartj/ehac244

6. Camilli M, Cipolla CM, Dent S, Minotti G, Cardinale DM. Anthracycline Cardiotoxicity in Adult Cancer Patients: JACC: CardioOncology State-of-the-Art Review. JACC CardioOncol 2024;6:655–677. doi: 10.1016/j.jaccao.2024.07.016

7. Fabiani I, Chianca M, Cipolla CM, Cardinale DM. Anthracycline-induced cardiomyopathy: risk prediction, prevention and treatment. Nat Rev Cardiol 2025;22:551–563. doi: 10.1038/s41569-025-01126-1

8. Lancellotti P, Suter TM, Lopez-Fernandez T, Galderisi M, Lyon AR, Van der Meer P, et al. Cardio-Oncology Services: rationale, organization, and implementation. Eur Heart J 2019;40:1756–1763. doi: 10.1093/eurheartj/ehy453

9. Ito-Hagiwara K, Hagiwara J, Endo Y, Becker LB, Hayashida K. Cardioprotective strategies against doxorubicin-induced cardiotoxicity: A review from standard therapies to emerging mitochondrial transplantation. Biomed Pharmacother 2025;189:118315. doi: 10.1016/j.biopha.2025.118315

10. Omland T, Heck SL, Gulati G. The Role of Cardioprotection in Cancer Therapy Cardiotoxicity: JACC: CardioOncology State-of-the-Art Review. JACC CardioOncol 2022;4:19–37. doi: 10.1016/j.jaccao.2022.01.101

11. Rawat PS, Jaiswal A, Khurana A, Bhatti JS, Navik U. Doxorubicin-induced cardiotoxicity: An update on the molecular mechanism and novel therapeutic strategies for effective management. Biomed Pharmacother 2021;139:111708. doi: 10.1016/j.biopha.2021.111708

12. Cardinale D, Colombo A, Lamantia G, Colombo N, Civelli M, De Giacomi G, et al. Anthracycline-induced cardiomyopathy: clinical relevance and response to pharmacologic therapy. J Am Coll Cardiol 2010;55:213–220. doi: 10.1016/j.jacc.2009.03.095

13. Cardinale D, Sandri MT, Colombo A, Colombo N, Boeri M, Lamantia G, et al. Prognostic value of troponin I in cardiac risk stratification of cancer patients undergoing high-dose chemotherapy. Circulation 2004;109:2749–2754. doi: 10.1161/01.CIR.0000130926.51766.CC

14. Serrano JM, Gonzalez I, Del Castillo S, Muniz J, Morales LJ, Moreno F, et al. Diastolic Dysfunction Following Anthracycline-Based Chemotherapy in Breast Cancer Patients: Incidence and Predictors. Oncologist 2015;20:864–872. doi: 10.1634/theoncologist.2014-0500

15. Stoodley PW, Richards DA, Boyd A, Hui R, Harnett PR, Meikle SR, et al. Altered left ventricular longitudinal diastolic function correlates with reduced systolic function immediately after anthracycline chemotherapy. Eur Heart J Cardiovasc Imaging 2013;14:228–234. doi: 10.1093/ehjci/jes139

16. Tassan-Mangina S, Codorean D, Metivier M, Costa B, Himberlin C, Jouannaud C, et al. Tissue Doppler imaging and conventional echocardiography after anthracycline treatment in adults: early and late alterations of left ventricular function during a prospective study. Eur J Echocardiogr 2006;7:141–146. doi: 10.1016/j.euje.2005.04.009

17. Upshaw JN, Finkelman B, Hubbard RA, Smith AM, Narayan HK, Arndt L, et al. Comprehensive Assessment of Changes in Left Ventricular Diastolic Function With Contemporary Breast Cancer Therapy. JACC Cardiovasc Imaging 2020;13:198–210. doi: 10.1016/j.jcmg.2019.07.018

18. Plana JC, Galderisi M, Barac A, Ewer MS, Ky B, Scherrer-Crosbie M, et al. Expert consensus for multimodality imaging evaluation of adult patients during and after cancer therapy: a report from the American Society of Echocardiography and the European Association of Cardiovascular Imaging. J Am Soc Echocardiogr 2014;27:911–939. doi: 10.1016/j.echo.2014.07.012

19. Koukorava C, Ahmed K, Almaghrabi S, Pointon A, Haddrick M, Cross MJ. Anticancer drugs and cardiotoxicity: the role of cardiomyocyte and non-cardiomyocyte cells. Front Cardiovasc Med 2024;11:1372817. doi: 10.3389/fcvm.2024.1372817

20. Wenningmann N, Knapp M, Ande A, Vaidya TR, Ait-Oudhia S. Insights into Doxorubicin-induced Cardiotoxicity: Molecular Mechanisms, Preventive Strategies, and Early Monitoring. Mol Pharmacol 2019;96:219–232. doi: 10.1124/mol.119.115725

21. Wallace KB, Sardao VA, Oliveira PJ. Mitochondrial Determinants of Doxorubicin-Induced Cardiomyopathy. Circ Res 2020;126:926–941. doi: 10.1161/CIRCRESAHA.119.314681

22. Asnani A, Moslehi JJ, Adhikari BB, Baik AH, Beyer AM, de Boer RA, et al. Preclinical Models of Cancer Therapy-Associated Cardiovascular Toxicity: A Scientific Statement From the American Heart Association. Circ Res 2021;129:e21–e34. doi: 10.1161/RES.0000000000000473

23. Podyacheva EY, Kushnareva EA, Karpov AA, Toropova YG. Analysis of Models of Doxorubicin-Induced Cardiomyopathy in Rats and Mice. A Modern View From the Perspective of the Pathophysiologist and the Clinician. Front Pharmacol 2021;12:670479. doi: 10.3389/fphar.2021.670479

24. Genet G, Guilbeau-Frugier C, Honton B, Dague E, Schneider MD, Coatrieux C, et al. Ephrin-B1 is a novel specific component of the lateral membrane of the cardiomyocyte and is essential for the stability of cardiac tissue architecture cohesion. Circ Res 2012;110:688–700. doi: 10.1161/CIRCRESAHA.111.262451

25. Kane LA, Lazarou M, Fogel AI, Li Y, Yamano K, Sarraf SA, et al. PINK1 phosphorylates ubiquitin to activate Parkin E3 ubiquitin ligase activity. J Cell Biol 2014;205:143–153. doi: 10.1083/jcb.201402104

26. Kondapalli C, Kazlauskaite A, Zhang N, Woodroof HI, Campbell DG, Gourlay R, et al. PINK1 is activated by mitochondrial membrane potential depolarization and stimulates Parkin E3 ligase activity by phosphorylating Serine 65. Open Biol 2012;2:120080. doi: 10.1098/rsob.120080

27. Liu JE, Barac A, Thavendiranathan P, Scherrer-Crosbie M. Strain Imaging in Cardio-Oncology. JACC CardioOncol 2020;2:677–689. doi: 10.1016/j.jaccao.2020.10.011

28. Oikonomou EK, Kokkinidis DG, Kampaktsis PN, Amir EA, Marwick TH, Gupta D, et al. Assessment of Prognostic Value of Left Ventricular Global Longitudinal Strain for Early Prediction of Chemotherapy-Induced Cardiotoxicity: A Systematic Review and Meta-analysis. JAMA Cardiol 2019;4:1007–1018. doi: 10.1001/jamacardio.2019.2952

29. Sartorio A, Cristin L, Pont CD, Farzaneh-Far A, Romano S. Global longitudinal strain as an early marker of cardiac damage after cardiotoxic medications, a state-of-the-art review. Prog Cardiovasc Dis 2025;89:92–101. doi: 10.1016/j.pcad.2025.01.001

30. Dean M, Kim MJ, Dimauro S, Tannenbaum S, Graham G, Liang BT, et al. Cardiac and noncardiac biomarkers in patients undergoing anthracycline chemotherapy - a prospective analysis. Cardiooncology 2023;9:23. doi: 10.1186/s40959-023-00174-1

31. Feng W, Wang Q, Tan Y, Qiao J, Liu Q, Yang B, et al. Early detection of anthracycline-induced cardiotoxicity. Clin Chim Acta 2025;565:120000. doi: 10.1016/j.cca.2024.120000

32. Galan-Arriola C, Lobo M, Vilchez-Tschischke JP, Lopez GJ, de Molina-Iracheta A, Perez-Martinez C, et al. Serial Magnetic Resonance Imaging to Identify Early Stages of Anthracycline-Induced Cardiotoxicity. J Am Coll Cardiol 2019;73:779–791. doi: 10.1016/j.jacc.2018.11.046

33. Mabudian L, Jordan JH, Bottinor W, Hundley WG. Cardiac MRI assessment of anthracycline-induced cardiotoxicity. Front Cardiovasc Med 2022;9:903719. doi: 10.3389/fcvm.2022.903719

34. Mohamed AA, Elmancy LY, Abulola SM, Al-Qattan SA, Mohamed Ibrahim MI, Maayah ZH. Assessment of Native Myocardial T1 Mapping for Early Detection of Anthracycline-Induced Cardiotoxicity in Patients with Cancer: a Systematic Review and Meta-analysis. Cardiovasc Toxicol 2024;24:563–575. doi: 10.1007/s12012-024-09866-1

35. Camilli M, Ferdinandy P, Salvatorelli E, Menna P, Minotti G. Anthracyclines, Diastolic Dysfunction and the road to Heart Failure in Cancer survivors: An untold story. Prog Cardiovasc Dis 2024;86:38–47. doi: 10.1016/j.pcad.2024.07.002

36. Minotti G, Reggiardo G, Camilli M, Salvatorelli E, Menna P. From Cardiac Anthracycline Accumulation to Real-Life Risk for Early Diastolic Dysfunction: A Translational Approach. JACC CardioOncol 2022;4:139–140. doi: 10.1016/j.jaccao.2021.12.002

37. Diaz-Guerra A, Villena-Gutierrez R, Clemente-Moragon A, Gomez M, Oliver E, Fernandez-Tocino M, et al. Anthracycline Cardiotoxicity Induces Progressive Changes in Myocardial Metabolism and Mitochondrial Quality Control: Novel Therapeutic Target. JACC CardioOncol 2024;6:217–232. doi: 10.1016/j.jaccao.2024.02.005

38. Guo X, Jiang M, Tao Z, Dai H, Wu C, Wang Y, et al. 5-Oxoproline prevents doxorubicin-induced cardiotoxicity and tumor growth. Redox Biol 2025;85:103753. doi: 10.1016/j.redox.2025.103753

39. Hamo CE, DeJong C, Hartshorne-Evans N, Lund LH, Shah SJ, Solomon S, et al. Heart failure with preserved ejection fraction. Nat Rev Dis Primers 2024;10:55. doi: 10.1038/s41572-024-00540-y

40. Piquereau J, Ventura-Clapier R. Maturation of Cardiac Energy Metabolism During Perinatal Development. Front Physiol 2018;9:959. doi: 10.3389/fphys.2018.00959

41. Bers DM. Cardiac excitation-contraction coupling. Nature 2002;415:198-205. doi: 10.1038/415198a

42. Shooshtarian AK, O’Gallagher K, Shah AM, Zhang M. SERCA2a dysfunction in the pathophysiology of heart failure with preserved ejection fraction: a direct role is yet to be established. Heart Fail Rev 2025;30:545–564. doi: 10.1007/s10741-025-10487-1

43. Zhihao L, Jingyu N, Lan L, Michael S, Rui G, Xiyun B, et al. SERCA2a: a key protein in the Ca(2+) cycle of the heart failure. Heart Fail Rev 2020;25:523–535. doi: 10.1007/s10741-019-09873-3

