## supplemental Figure 1-5 for "Impaired active myocardial relaxation is the earliest sign of anthracycline cardiotoxicity and reflects selective injury of the cardiomyocyte surface"

### Supplementary Figures

### Supplemental Figure 1

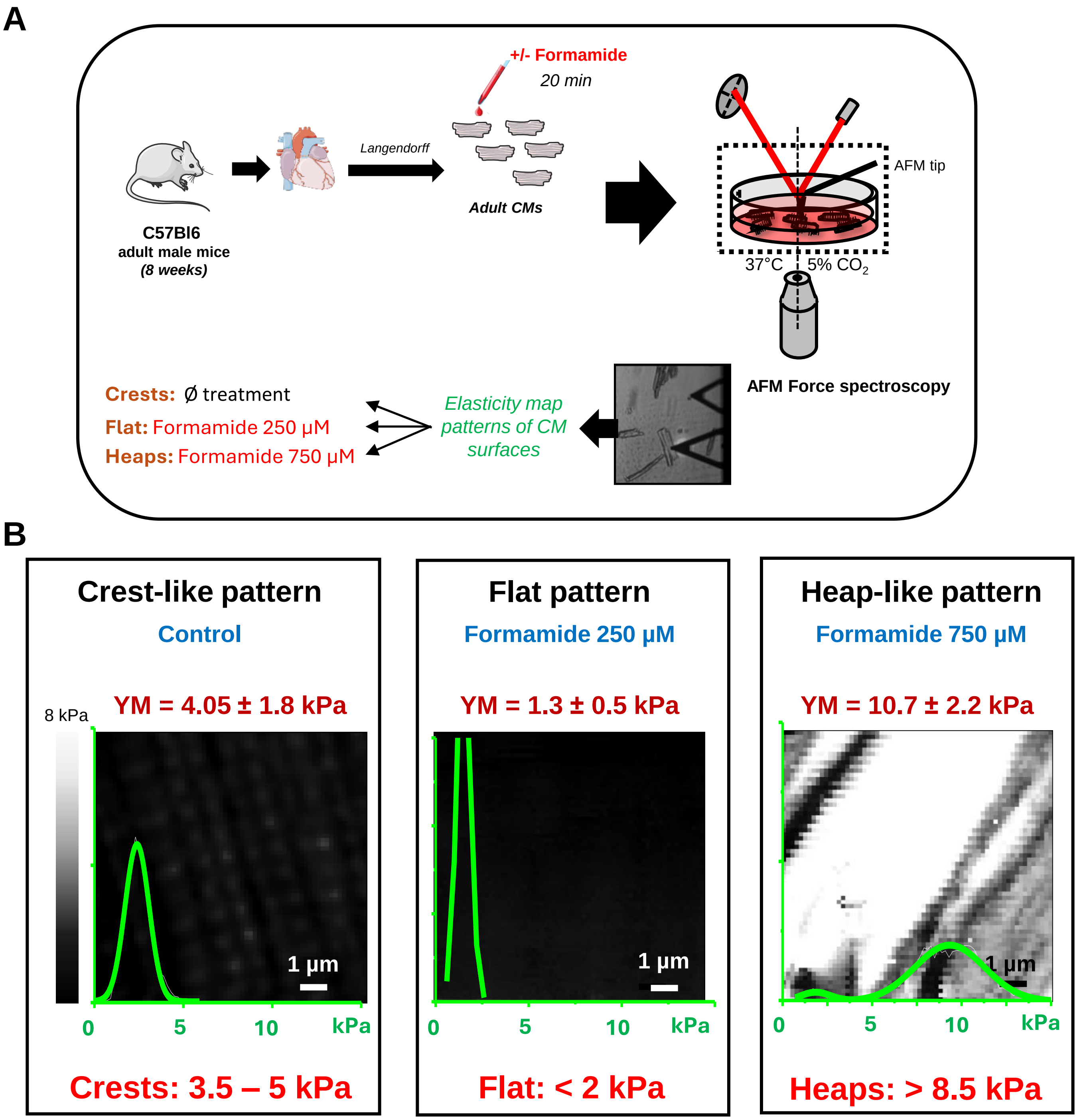


**Supplemental Figure 1. Definition of cardiomyocyte (CM) surface patterns by their elasticity signatures.**

(**A**)Schematic representation of the experimental protocol. Freshly isolated cardiomyocytes (CMs) from adult male C57BL/6J mice (8 weeks) were obtained by Langendorff perfusion and analyzed by atomic force microscopy (AFM) in force-volume mode (37°C, 5% $CO_2$). To reproduce the three CM surface patterns previously identified *in vivo* and validate their detection on a new AFM setup, isolated CMs were exposed for 20 min to increasing formamide concentrations, as previously described[2]. Each concentration was selected for its ability to reliably induce a specific pattern: no formamide (crest-like), 250 µM (flat) and 750 µM (heap-like).

(**B**) Representative elasticity maps (64 × 64 force curves over a 100 µm² area) illustrate the three main CM surface patterns and their corresponding Young's modulus (YM): Crest-like pattern (control, untreated CMs): organized surface topography and intermediate stiffness (YM = 4.05 ± 1.8 kPa; range: 3.5-5 kPa); Flat pattern (Formamide 250 µM): reduced topographical complexity and decreased stiffness (YM = 1.3 ± 0.5 kPa; YM < 2 kPa); Heap-like pattern (Formamide 750 µM): prominent surface protrusions and increased stiffness (YM = 10.7 ± 2.2 kPa; YM > 8.5 kPa). n = 1 mouse, 3 CMs, 12,288 force curves per condition. Scale bars, 2 µm. Data are mean ± SD.

### Supplemental Figure 2

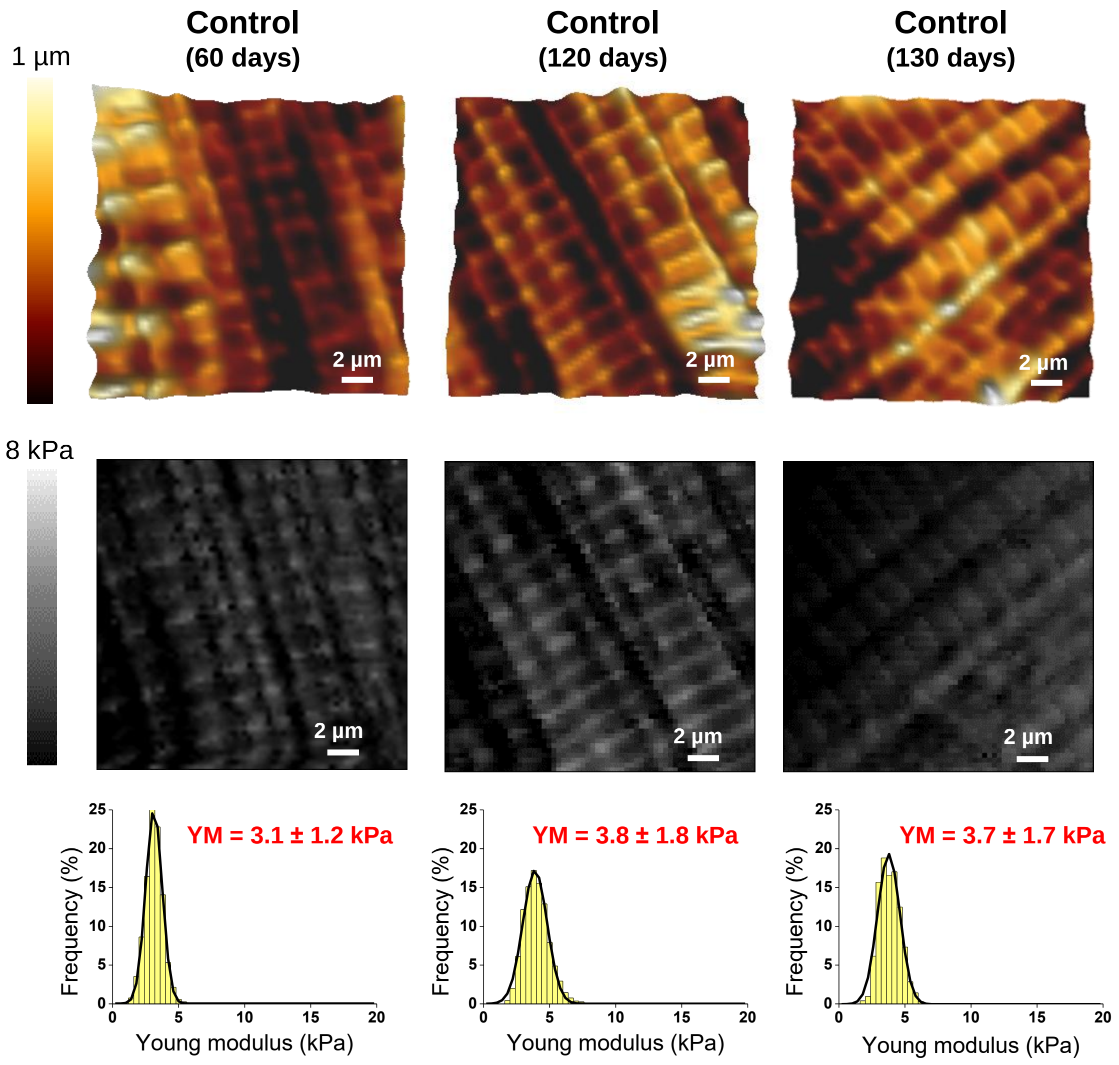


**Supplemental Figure 2. Cardiomyocyte surface architecture and stiffness are stable over time under control conditions.**

Freshly isolated cardiomyocytes (CMs) from saline-injected adult male C57BL/6J mice (8 weeks old) were analyzed by atomic force microscopy (AFM) in force-volume mode at successive time points (60, 120 and 130 days).

(**Top**) Representative AFM topography images (10 × 10 µm scan area) showing preserved surface morphology across time points

(**Middle**) Corresponding elasticity maps (4,096 force curves/cell; 10×10 µm) showing a homogeneous elasticity distribution.

(**Bottom**) Frequency distribution of Young's modulus (YM) from all force curves, with Gaussian fitting to determine the peak YM/group (red). Surface stiffness remained stable across time points (YM≈3-4 kPa; crest-like range). n= 61,440 curves/time point; 12-15 CMs from 4-5 mice/condition. Data are mean ± SD.

### Supplemental Figure 3

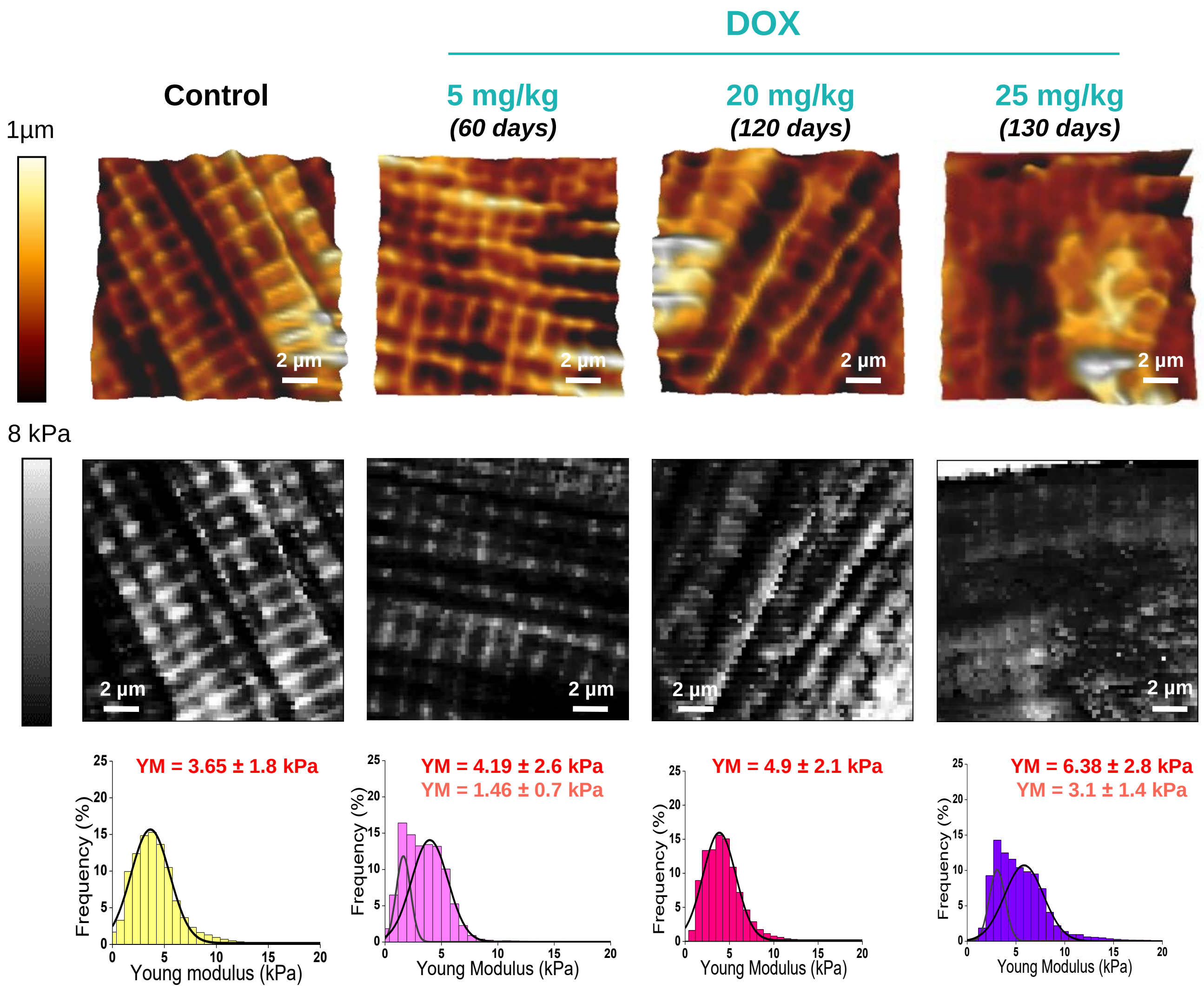


**Supplemental Figure 3. Qualitative and quantitative assessment of cardiomyocyte surface stiffness remodeling following *in vivo* doxorubicin treatment.**

Freshly isolated adult CMs from mice exposed to increasing cumulative DOX doses (5 to 25 mg/kg) were analyzed using atomic force microscopy (AFM) in Force Spectroscopy mode. For each condition, three levels of analysis are shown: (*top*) Representative surface topography (10 µm × 10 µm); (*middle*) Elasticity map (10 µm × 10 µm; 4096 force curves); (*bottom*) Frequency distribution of Young's modulus (YM) extracted from individual force curves. Gaussian models were applied to each distribution to extract peak YM values.

Control CMs (saline-treated, combined day 60-120-130) exhibit a unimodal distribution centered at 3.65 ± 1.8 kPa (n=50 cells, 18 mice, 209,800 curves). Following low-dose DOX exposure (5 mg/kg, day 60), the YM distribution shifted to a bimodal profile, with distinct peaks at 1.46 ± 0.7 kPa and 4.19 ± 2.6 kPa (n=21 cells, 5 mice, 86,016 curves), indicative of surface softening in a subset of CMs. At a higher cumulative dose (25 mg/kg, day 130), YM values increased, yielding a broader unimodal distribution centered at 4.9 ± 2.1 kPa (n=20 cells, 5 mice, 81,920 curves), suggesting progressive stiffening consistent with IFM aggregation at the cell surface.

### Supplemental Figure 4

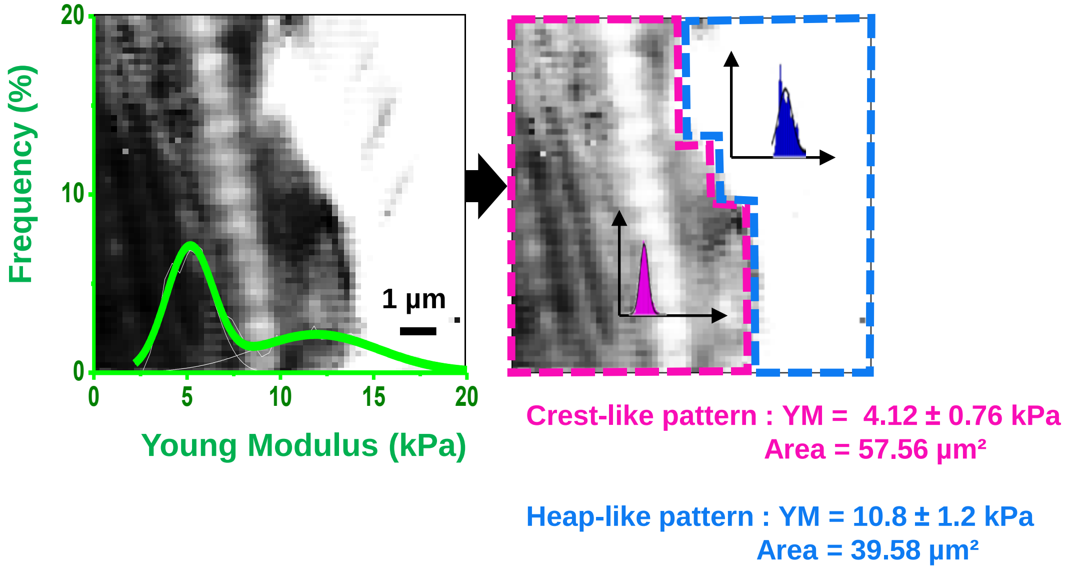


| Experimental group | Crest-like pattern | | Flat pattern | | Heap-like pattern | |
|---|---|---|---|---|---|---|
| | YM (kPa) | Area (µm²) | YM (kPa) | Area (µm²) | YM (kPa) | Area (µm²) |
| 25 mg/kg | 4.12 | 57.56 | 0 | 0 | 10.8 | 39.58 |

**Supplemental Figure 4. Strategy for analyzing heterogeneous cardiomyocyte surface elasticity maps using motif-specific segmentation.**
(**Left**) Representative elasticity map (10 × 10 µm²) from a CM isolated 36 days after a cumulative DOX dose of 30 mg/kg, showing a bimodal Young's modulus (YM) distribution overlaid on the image (green line). Each peak in the YM distribution corresponds to a distinct surface motif previously defined in Figure S1. Based on color-coded elasticity ranges, regions matching crest-like and heap-like motifs were segmented and analyzed separately.
(**Right**) Schematic of the segmentation approach applied to each elasticity map, assigning pixels to one of three predefined motifs (crest, flat, or heap) based on YM thresholds and topographical features. For each motif, mean elasticity (kPa) and corresponding surface area (µm²) were computed and recorded in a structured results table (excerpt shown at bottom). In the example shown, the analyzed map revealed coexisting crest and heap motifs with average elasticities of 4.12 ± 0.76 kPa and 10.8 ± 1.2 kPa, and corresponding surface areas of 57.56 µm² and 39.58 µm², respectively. This motif-specific analysis was systematically applied across all conditions to quantify spatially heterogeneous mechanical remodeling of the CM surface during DIC progression.

### Supplemental Figure 5

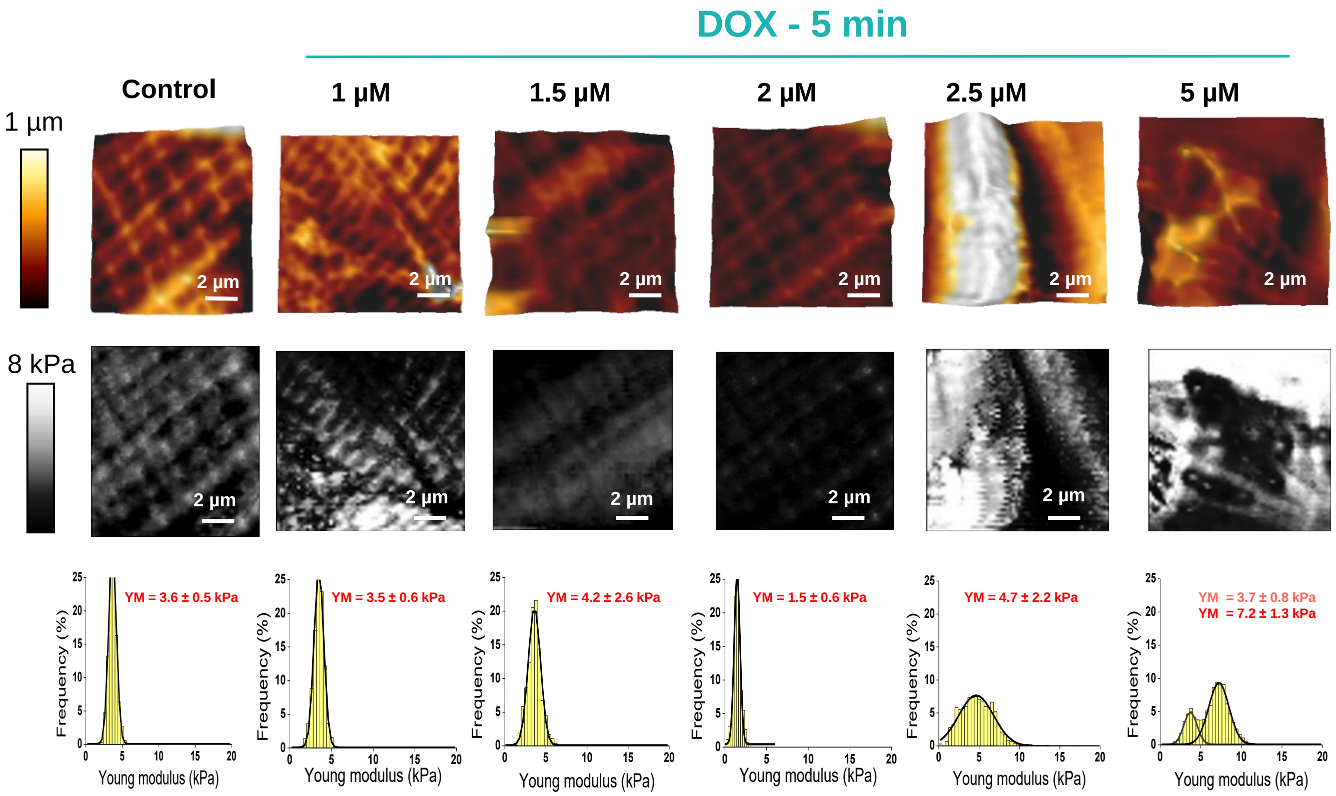


**Supplemental Figure 5. *In vitro* exposure to doxorubicin induces dose-dependent alterations in cardiomyocyte surface stiffness.**

Freshly isolated CMs from adult male C57BL/6J mice (8 weeks old) CMs were incubated for 5 minutes at 37°C with increasing concentrations of DOX (1-5 µM). Cell surface mechanical properties were assessed using AFM in Force Spectroscopy mode over a 10 × 10 µm² area (64 × 64 force curves; total of 36,864 curves/condition; n = 9-19 cells; 3 mice/group).

**(Top)** Representative 3D surface topography images for each condition.

**(Middle)** Corresponding elasticity maps showing spatial distribution of Young's modulus (YM).

**(Bottom)** Frequency histograms of YM values extracted from AFM force curves. Gaussian curve fitting was used to determine the predominant YM values. Control CMs (untreated) exhibited a unimodal distribution centered at 3.6 ± 0.5 kPa; At 1.5 µM DOX, YM increased to 4.2 ± 2.6 kPa; At 2 µM DOX, YM shifted toward lower stiffness (1.5 ± 0.6 kPa); At 2.5 µM DOX, YM increased to 4.7 ± 2.2 kPa; At 5 µM DOX, the distribution became bimodal, with peaks at 3.7 ± 0.8 kPa and 7.2 ± 1.3 kPa, indicating the emergence of localized stiffened subdomains on the CM surface.
